# Deficiency of the splicing factor RBM10 limits EGFR inhibitor response in *EGFR* mutant lung cancer

**DOI:** 10.1101/2020.10.26.356352

**Authors:** Shigeki Nanjo, Wei Wu, Niki Karachaliou, Collin M. Blakely, Junji Suzuki, Siraj Ali, D. Lucas Kerr, Victor Olivas, Jonathan Shue, Julia Rotow, Manasi Mayekar, Franziska Haderk, Nilanjana Chatterjee, Anatoly Urisman, Yuriy Kirichok, Daniel S. W. Tan, Rafael Rosell, Ross A Okimoto, Trever G. Bivona

## Abstract

Molecularly targeted cancer therapy has improved outcomes for cancer patients with targetable oncoproteins, such as mutant *epidermal growth factor receptor* (*EGFR*) in lung cancer. Yet, long-term patient survival remains limited because treatment responses are typically incomplete. One potential explanation for the lack of complete and durable responses is that oncogene-driven cancers with activating mutations in the *EGFR* often harbor additional co-occurring genetic alterations. This hypothesis remains untested for most genetic alterations that co-occur with mutant *EGFR*. Here, we report the functional impact of inactivating genetic alteration of the mRNA splicing factor *RBM10* that co-occur with mutant *EGFR*. RBM10 deficiency decreased EGFR inhibitor efficacy in patient-derived *EGFR* mutant tumor models. RBM10 modulated mRNA alternative splicing of the mitochondrial apoptotic regulator Bcl-x to regulate tumor cell apoptosis during treatment. Genetic inactivation of *RBM10* diminished EGFR inhibitor-mediated apoptosis by decreasing the ratio of Bcl-xS-(pro-apoptotic)-to-Bcl-xL(anti-apoptotic) Bcl-x isoforms. RBM10 deficiency was a biomarker of poor response to EGFR inhibitor treatment in clinical samples. Co-inhibition of Bcl-xL and mutant EGFR overcame resistance induced by RBM10 deficiency. This study sheds light on the role of co-occurring genetic alterations, and on the impact of splicing factor deficiency in the modulation of sensitivity to targeted kinase inhibitor cancer therapy.

## Introduction

Lung cancer is the leading cause of cancer mortality among both men and women, comprising almost 25% of all cancer deaths (1). There has been significant progress in the treatment of lung cancer, and many other cancer types, in the last 10 years through the advent of precision medicine that leverages tumor molecular and genetic profiling coupled with molecularly targeted cancer therapy. As one paradigm-defining example of precision medicine, activating mutations in the *Epidermal Growth Factor Receptor* (*EGFR*) are associated with high response rates to EGFR-directed tyrosine kinase inhibitors (TKIs) in non-small cell lung cancer (NSCLC) (1). The most common TKI sensitizing mutations in the *EGFR* are deletions in *exon 19* that affect the LREA motif and substitutions in *exon 21* (*L858R*), which together account for >90% of all *EGFR* mutations in lung adenocarcinoma (LA) (2). While these canonical *EGFR* mutations typically confer sensitivity to EGFR TKIs, approximately 20-30% of patients exhibit either primary refractory disease (intrinsic resistance), or a limited response (such as less than 30% tumor regression by Response Evaluation Criteria in Solid Tumors (RECIST) version1.1 criteria (3)) followed by disease progression (4–7). While several mechanisms of intrinsic resistance have been reported (8–12), the mechanisms underlying the distinct clinical scenario of limited tumor response followed by earlier tumor progression during initial EGFR TKI treatment are less well defined. This is perhaps highlighted through the clinical observation that EGFR mutant patients that harbor the *EGFR L858R* mutation experience shorter progression free survival (PFS) to identical EGFR inhibitor treatment compared to patients with *EGFR exon 19 deletions* in most clinical trials for reasons that are not well understood (4–7).

The use of next generation sequencing (NGS) platforms that profile large panels of cancer-relevant genes has shown that *EGFR* mutant tumors often harbor additional co-occurring genetic alterations, both before and following therapy (13–15). One hypothesis arising from this observation is that certain co-occurring alterations present in an *EGFR* mutant tumor could modulate sensitivity to EGFR TKI treatment and explain, in part, the variable magnitude and duration of anti-tumor treatment response in patients. With a few important exceptions such as genetic alterations in *TP53* (13, 14), the functional impact (if any) of most of these co-occurring genetic alterations on tumor growth or treatment sensitivity is not well established.

One mechanism by which co-occurring genetic alterations could impact therapeutic sensitivity to EGFR TKI treatment is via the modulation of tumor cell apoptosis (16). The Bcl-2 family of proteins regulates the intrinsic pathway of mitochondrial-mediated apoptosis (17) and comprises both pro-apoptotic and anti-apoptotic components. The pro-apoptotic arm of this pathway is often activated by cancer treatment, which initiates the depolarization of the mitochondrial outer membrane potential and release of cytochrome c into the cytoplasm to form the apoptosome, which subsequently activates Caspase-3 and Caspase-7 that are effector caspases (18). In *EGFR* mutant tumors, up-regulation of the pro-apoptotic Bcl-2 family protein BIM is an important event that is required for EGFR inhibitor-induced apoptosis (16). Genetic loss of *BIM* can limit EGFR inhibitor therapeutic response (11, 12). Beyond BIM modulation, how *EGFR* mutant tumor cells regulate the apoptotic response during EGFR targeted therapy remains incompletely defined.

mRNA alternative splicing is one mechanism used by cells to generate phenotypic and functional diversity that impacts on a range of cellular behaviors (19, 20). Recent studies identified a role for mRNA splicing factors in oncogenesis in hematologic malignancies (21, 22). Recent reports also indicate the presence of genetic alterations in mRNA splicing genes in solid malignancies (23). For instance, the Cancer Genome Atlas (TCGA) lung adenocarcinoma (LA) profiling analysis showed mutation of the splicing factor *RBM10* (*RNA binding motif 10*) at a relatively high frequency (24, 25). The potential role of these RBM10 genetic alterations in LA is not well defined. RBM10 can regulate mRNA alternative splicing and may act as a tumor suppressor in some contexts (26–28). Here, we investigated the open question of whether, and how mutations in splicing factors such as *RBM10* contribute to lung cancer pathogenesis or modulate sensitivity to oncoprotein targeted kinase inhibitor therapy.

## Results

### Identification of potential genetic modifiers of EGFR inhibitor response highlights *RBM10*

Oncogenic *EGFR* activating mutations are associated with high initial response rates to EGFR-directed therapy in human LA (1). However, 20-30% of patients with *EGFR* mutations will either not respond with demonstrable (i.e. >30%) tumor regression or develop relatively early tumor progression following an initial incomplete response (3) to EGFR TKI treatment (4–7). Tumor genetic heterogeneity that is present prior to therapy could contribute to this incomplete initial response or early emergence of resistance. In order to identify genetic modifiers of therapeutic response in *EGFR* mutant LA, we performed targeted next generation sequencing (NGS) of 324 cancer-relevant genes in 591 *EGFR* mutant LA human tumors (median coverage depth = 500x) (see Methods). Through this analysis, we noted frequent truncating mutations in the *RBM10* gene (7.6%) (Figure 1A). A similar frequency of *RBM10* truncating mutations was present in the MSK-IMPACT *EGFR* mutant LA dataset (8.0%) (Supplementary Figure 1A) (29). Interestingly, we observed that 92% of *RBM10* alterations present in our *EGFR* mutant tumor dataset did not co-occur with a known EGFR TKI resistance-associated mutation such as *EGFR T790M* or *C797S* or *MET* gene amplification (Supplementary Figure 1B). This suggests a potential modifying role for *RBM10* mutations in these *EGFR* mutant tumors outside of known genetic mechanisms of resistance (Figure 1B). Consistent with prior observations, *RBM10* mutations in our dataset primarily resulted in premature truncation of the protein coding sequence, suggesting a loss-of-function phenotype (Figure 1C) (23–25). In general, *RBM10* mutations were more often subclonal compared to the founder *EGFR* mutation (Fisher exact test p=0.02, Supplementary Figure 1C) in *EGFR* mutant tumors (30). Since there is no known functional impact of *RBM10* in *EGFR* mutant lung cancer treatment, we tested the hypothesis that *RBM10* inactivation modulates EGFR TKI sensitivity in *EGFR* mutant LA.

**Figure 1.**
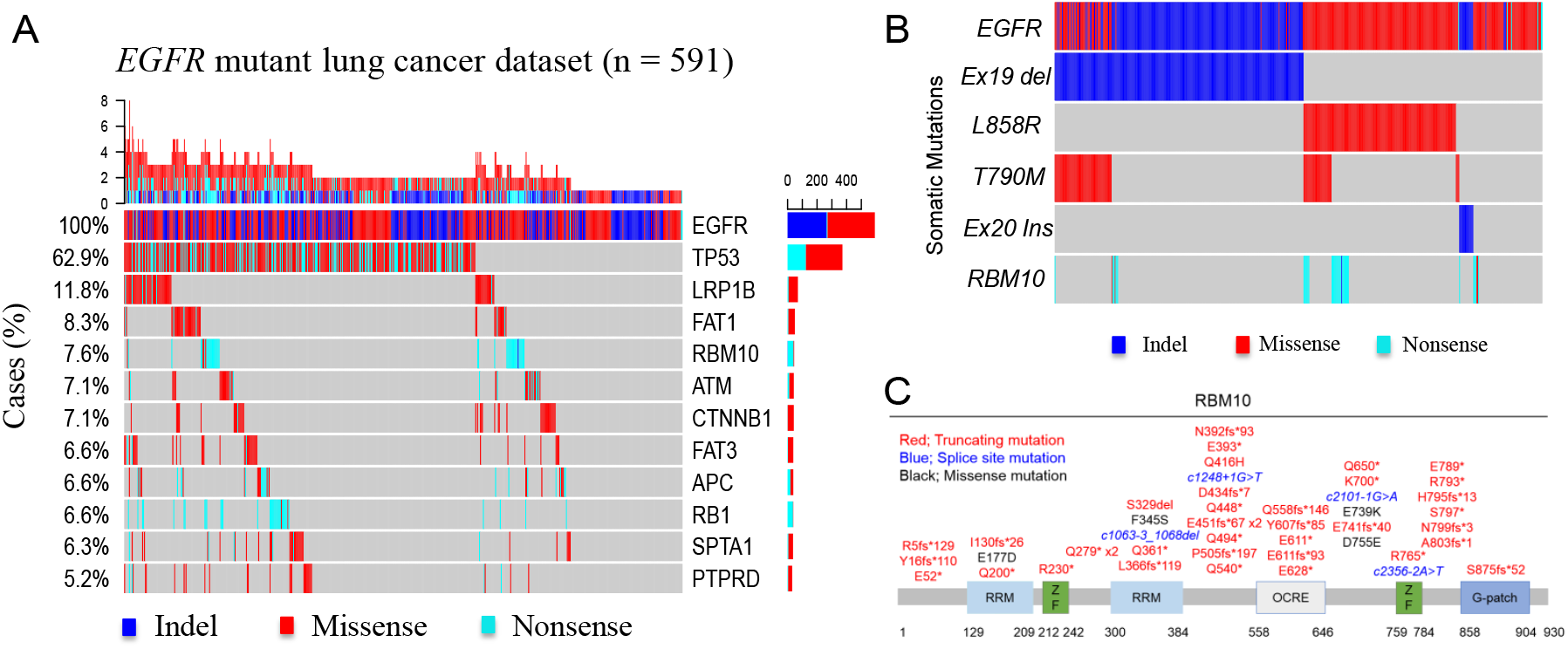
*RBM10* mutations co-occur with *EGFR* mutations in lung adenocarcinoma. **A,** Targeted next generation sequencing (NGS) of 591 *EGFR* mutant lung adenocarcinoma (LA) tumors using a panel of 324 cancer related genes (median coverage depth = 500x). Co-occurring alterations that occurred in at least 5% of *EGFR* mutation positive cases are shown. **B,** *RBM10* alterations were compared across each *EGFR* mutant subtype. **C,** Mutations in the *RBM10* protein coding sequence (Splice site mutations: blue, Truncating mutations: red, Missense mutations: black). RRM; RNA recognition motifs, ZF; Zinc finger, G-patch; glycine-patch, OCRE; octamer repeat sequence.

To test this hypothesis, first we performed functional genetic experiments by *RBM10* knockdown (KD) and overexpression (OE) in patient-derived lung cancer cell lines to assess cell viability and apoptotic response to the EGFR kinase inhibitor osimertinib, which we used in clinically-relevant concentrations (31). Increased cell viability during osimertinib treatment was observed upon genetic silencing of *RBM10* with two independent shRNAs in patient-derived LA cell lines H3255 (*EGFR L858R*) and PC-9 (*EGFR del19*) compared to control (Supplementary Figure 2A-F). To assess whether RBM10 expression could modulate apoptosis in *EGFR* mutant lung cancer, we first assessed cleavage of the apoptotic biomarker PARP in H3255 (*EGFR L858R/RBM10* WT) and PC-9 (*EGFR del19*/*RBM10* WT) cells. Indeed, after osimertinib treatment cleaved-PARP levels were decreased in *RBM10* KD cells compared to control (Figure 2A-B). Additionally, the activities of the key apoptotic effector Caspases-3 and −7 were similarly decreased upon *RBM10* KD in H3255 and PC-9 cells treated with osimertinib (Figure 2C-D). We next studied a recently established *EGFR L858R* mutant patient-derived cell line (A014) that harbors a co-occurring *RBM10* truncating mutation and displays relative resistance to EGFR inhibitors, showing minimal apoptosis upon EGFR TKI treatment (Figure 2E-F) (15). We reconstituted wild type (WT) *RBM10* into these RBM10-deficient A014 cells and observed enhanced apoptosis upon osimertinib treatment, as measured by cleaved PARP levels and Caspase 3 and 7 activity (Figure 2E-F). These data highlight the RBM10 functional deficiency that characterizes the *RBM10* mutant A014 patient-derived model. In the absence of osimertinib treatment, *RBM10* KD did not initiate tumor formation in non-cancerous Beas2B human bronchial epithelial cells lacking canonical driver mutations (Supplementary Figure 3A) nor did *RBM10* KD significantly enhance tumor growth and proliferation in H3255 or PC-9 patient-derived *EGFR* mutant LA *in vivo* subcutaneous tumor models (Supplementary Figure 3C-F). Collectively, these findings suggest that RBM10 expression impacts the therapeutic sensitivity to EGFR targeted therapy by modulating the apoptotic response, without a substantial impact on tumor cell proliferation or on tumor initiation or growth in the absence of either oncogenic *EGFR* or EGFR inhibitor treatment in these various systems.

**Figure 2.**
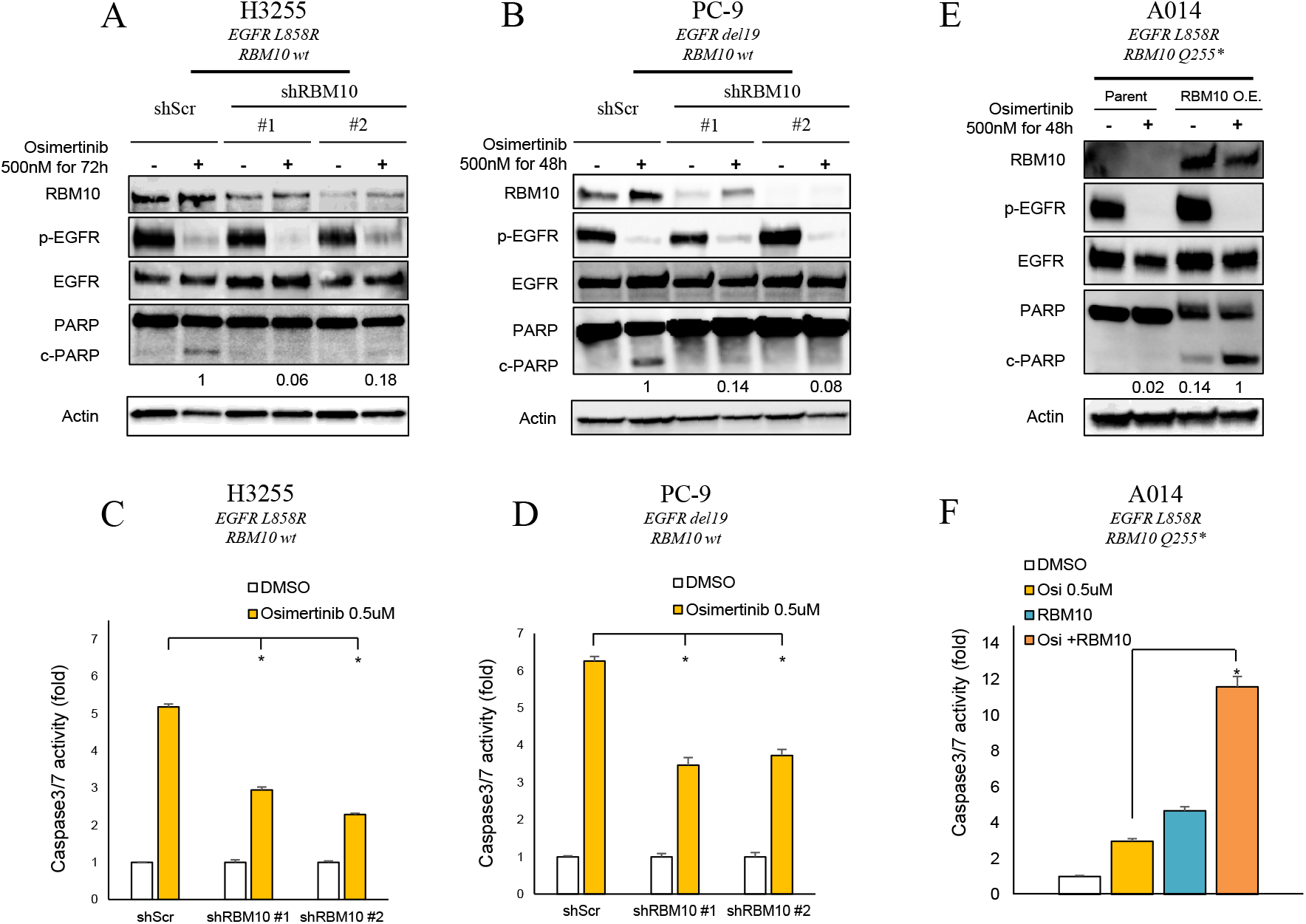
RBM10 modulates the apoptotic response to osimertinib in *EGFR* mutant lung adenocarcinoma. **A-D,** H3255 and PC-9 (*EGFR* mutant and *RBM10* wt) cells expressing *shRBM10* or shScramble control were treated with the third-generation EGFR inhibitor osimertinib (500 nM) or DMSO for 48 to 72 hours. Western blot analysis of the indicated proteins from cellular protein extracts was normalized to actin; quantification of cleaved PARP was determined by signal densitometry (**A, B**). Apoptotic response was assessed using a Caspase-Glo 3/7 assay. Each bar represents the mean ± SEM of the fold change after normalization to DMSO control. (**C, D**). **E-F,** RBM10-deficient A014 (*EGFR* mutant and *RBM10 Q255*\*) cells with genetic reconstitution of WT *RBM10* were treated with osimertinib (500 nM) for 48 hours. Western blotting of indicated lysates was normalized to actin (**E**). Caspase-3/7 activity was measured using a Caspase-Glo 3/7 assay. Each bar represents the mean ± SEM of the fold change after normalization to DMSO control. (**F**). Data represent 3 independent experiments. O.E.:Overexpression. *; p < 0.05.

We next tested if co-occurring *RBM10* inactivation in *EGFR* mutant tumors contributes to EGFR inhibitor resistance *in vivo*. We treated mice bearing H3255 or PC-9 tumor xenografts with osimertinib or vehicle and found that mice with tumors in which *RBM10* was silenced had decreased osimertinib sensitivity compared to shScramble control (Figure 3A-B and Supplementary Figure 3C-D). Using clinical response criteria (32), we observed that *EGFR* mutant tumors harboring sh*RBM10* had decreased depth and frequency of response (Figure 3C-D). Consistent with our *in vitro* findings, we found decreased PARP cleavage in osimertinib treated tumors explanted from mice bearing tumors in which *RBM10* was silenced compared to control (Figure 3E-F and Supplementary Figure 4A-B).

**Figure 3.**
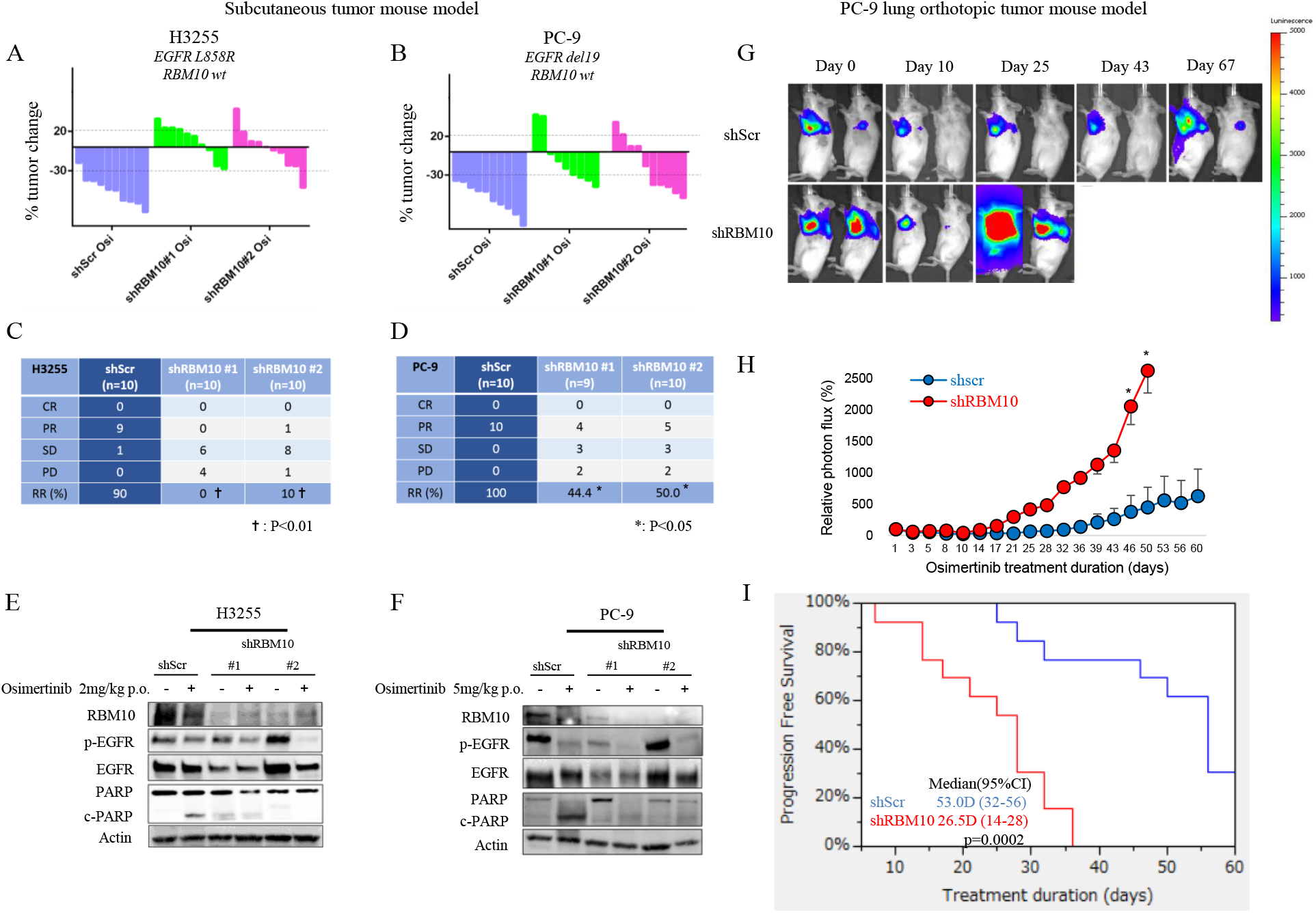
RBM10 deficiency limits the therapeutic efficacy of EGFR TKIs. **A-D,** Waterfall plots representing immunodeficient mice bearing H3255 (**A, C**) or PC-9 (**B, D**) tumor xenografts expressing either shScramble (shScr) control or sh*RBM10*; mice were treated with 2 mg/kg (H3255) or 5mg/kg (PC-9) osimertinib once daily over a duration of 14 days (n=10 tumors per treatment cohort). Percent changes in tumor volume compared to baseline volume (Day 0) for individual tumor xenografts are shown (**A, B**). Objective tumor response was graded using RECIST response criteria, comparing H3255 and PC-9 tumors expressing either shScr control or *shRBM10* treated with osimertinib (**C, D**). **E-F,** H3255 and PC-9 tumor xenograft explants demonstrating the effect of *RBM10* knockdown on PARP cleavage in mice treated with osimertinib or vehicle for 14 days. Tumors were harvested 4 hours after indicated treatments, and subsequent analyses of the indicated proteins was performed by western blot. **G-I,** PC-9 cells expressing either shScr or *shRBM10* in a validated orthotopic lung tumor model were treated with 5 mg/kg osimertinib once daily for 60 days. Representative bioluminescent images (**G**) and mean relative photon flux (**H**) are shown. Progression-free survival (PFS) comparing PC-9 shScr control and PC-9 *shRBM10* mice is shown (**I**) (p-value = 0.0002, Wilcoxon test). *; p < 0.05, †; p <0.01.

In order to model PFS in *EGFR* mutant LA, we leveraged our experience with an *in vivo* luciferase-based orthotopic lung cancer model (33, 34). We surgically implanted luciferase-labeled PC-9 cells expressing either scramble control or sh*RBM10* and treated these mice with osimertinib. We observed earlier initial tumor progression in mice bearing *RBM10* KD tumors compared to control in osimertinib-treated mice (Figure 3G-I) (Wilcoxon test p-value = 0.0002, n=13 each group). Thus, RBM10 suppression limits the initial response to EGFR targeted therapy in multiple patient-derived *in vitro* and *in vivo* systems.

### Clinical impact of RBM10 downregulation in advanced-stage *EGFR* mutant lung cancer treated with an EGFR TKI

Our preclinical data showed that low levels of RBM10 limited the response to EGFR TKI treatment. Therefore, we investigated the relationship between RBM10 expression levels and EGFR TKI treatment response in human advanced-stage (IIIB/IV) *EGFR* mutant NSCLC. We performed quantitative real-time PCR (QRT-PCR) analysis of *RBM10* in a panel (n=57) of clinically annotated *EGFR* mutant tumors obtained from patients treated with EGFR inhibitors. While we were unable to perform direct genomic sequencing of *RBM10* in this clinical cohort, we and others have observed a significant association between *RBM10* mRNA expression and *RBM10* mutation status, wherein *RBM10* mutations are associated with decreased *RBM10* mRNA expression (Supplementary Figure 4C) (35). We stratified *EGFR* mutant patients by median *RBM10* mRNA expression and found that patients whose tumors expressed lower levels of *RBM10* progressed earlier on EGFR TKI therapy (PFS: *RBM10* low = 11.6 months, *RBM10* high = 16.7 months, Wilcoxon test p-value = 0.03) (Figure 4A). Decreased PFS in the *RBM10* low cohort was associated with decreased initial tumor response rates (Figure 4B). Our *in vivo* mouse model findings (Figure 3) coupled with these clinical observations suggest that loss of *RBM10* can limit the initial response to EGFR targeted therapy by suppressing tumor cell apoptosis, leading to worse clinical outcomes in *EGFR* mutant LA patients.

**Figure 4.**
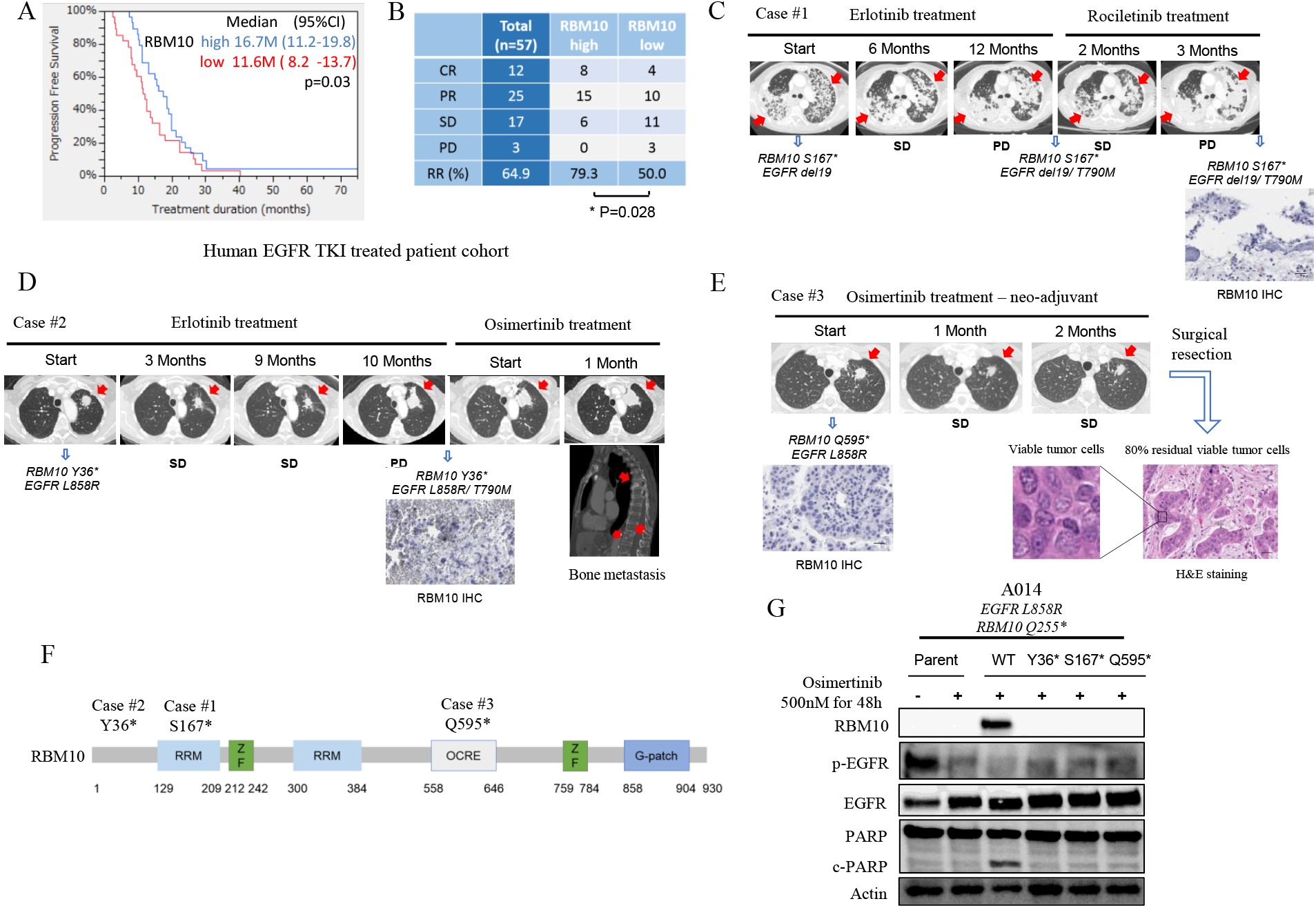
RBM10 deficiency is a biomarker of worse EGFR TKI response in human *EGFR* mutant lung cancer. **A-B,** A human EGFR TKI-treated patient cohort (n=57) was stratified into RBM10 high and low mRNA expression cohorts. PFS (p-value, Wilcoxon test) and response rate (p-value, Student t-test) in RBM10 high and low expression cohorts are shown. **C-E,** Somatic alterations were detected by NGS panel analysis of the tumor DNA from the patients. Case #1 harbored co-occurring mutations in *EGFR del19* and *RBM10 S167\** prior to EGFR inhibitor treatment. This patient had stable disease (SD) on erlotinib therapy (6 months), followed by early progression on the third-generation EGFR TKI rociletinib with the acquisition of an *EGFR T790M* mutation (**C**). Case #2 had co-occurring *EGFR L858R* and *RBM10 Y36\** mutations and demonstrated SD during erlotinib treatment (10 months). Following progression on erlotinib, the patient had progressive local and metastatic disease on osimertinib (**D**). Case #3 was a patient enrolled in a neodjuvant osimertinib clinical trial and found to harbor co-occurring *EGFR L858R* and *RBM10 Q595*\* mutations. Following two months of osimertinib treatment, radiographic measurements indicated SD and pathologic evaluation of the resected tumor specimen showed 80% viable tumor cells by H&E (hematoxylin and eosin) staining. RBM10 protein expression by immunohistochemistry (IHC) in patient-derived specimens obtained at either the time of progression (Case #1 and #2) or pre-neoadjuvant (Case #3) EGFR TKI therapy are shown at 200X magnification. Scale bars, 100 μm in (**C-E**). **F,** Engineered *RBM10 Y36*, RBM10 S167\**, and *RBM10 Q595\** mutations and the epitope sequences of antibodies used are shown. **G,** Immunoblot analysis of A014 (*RBM10 Q255*\*) cells transfected with constructs overexpressing *RBM10* WT or mutant forms (*Y36*, S167*, Q595*\*). Cells were treated with osimertinib (500 nM) or DMSO for 48 hours and western blot analysis was performed on cellular extracts. Data represent 3 independent experiments.

### Clinico-genomic validation of mutant *RBM10* in *EGFR* mutant lung cancer

We further assessed the clinical relevance of *RBM10* mutations in patients. We queried a distinct UCSF-based cohort to identify *EGFR* mutant lung cancer patients whose tumors harbored co-occurring *RBM10* mutations. We reasoned that loss-of-function genetic alterations in *RBM10* that dampen the apoptotic response to EGFR inhibitor therapy could enhance cancer cell survival in patients with *EGFR* mutant lung cancer. We confirmed the truncating mutations in the UCSF cohort decreased RBM10 protein expression by immunohistochemistry (IHC) in patient-derived specimens obtained at either the time of progression on EGFR TKI treatment (Case #1 and #2) or following neoadjuvant EGFR TKI treatment (Case #3; NCT03433469). Case #1 harbored the co-occurring mutations *EGFR del19* and *RBM10 S167*\* prior to EGFR inhibitor treatment. This patient’s tumor showed only stable disease (SD) on erlotinib therapy (6 months), followed by early progression that coincided with the acquisition of the drug resistant, *T790M* mutation in *EGFR* (Figure 4C). In a second patient (Case #2) with co-occurring *EGFR L858R* and *RBM10 Y36*\* mutations, again only SD was observed during erlotinib treatment (10 months) (Figure 4D). Biopsy at the time of progression confirmed low RBM10 expression (Figure 4D). Osimertinib was initiated in this case and there was immediate disease progression despite this treatment (Figure 4D). These clinical data are consistent with most of our experimental findings showing that genetic inactivation of *RBM10* limits the initial apoptotic response in *EGFR* mutant tumor cells. In certain cases (e.g. Case #1), RBM10 loss may mediate escape from tumor cell apoptosis during initial treatment and before the subsequent emergence of acquired resistance mutations such as *EGFR T790M* that drive tumor cell proliferation.

To associate radiographic response with pathological response, we next followed a patient (Case #3) treated with neoadjuvant osimertinib on a clinical trial (NCT03433469) whose tumor harbored the *EGFR L858R* and *RBM10 Q595\** co-occurring mutations. Following two months of neo-adjuvant osimertinib treatment, we confirmed low tumor cell RBM10 expression and observed radiographic SD, with 80% tumor cell viability in the osimertinib-treated, resected tumor specimen upon clinical pathologist assessment (Figure 4E). In a fourth clinical case, an *EGFR L858R* NSCLC harbored a co-occurring *RBM10 c2167-1 G>T* splice site mutation as detected by clinical-grade NGS. This patient was enrolled in the same neo-adjuvant osimertinib clinical trial and also showed minimal EGFR TKI response, with 68.3% tumor cell viability in the osimertinib-treated, resected tumor specimen upon clinical pathologist assessment (Supplementary Figure 5A, B). Consistent with these clinical data, other investigators observed early clinical progression associated with decreased sensitivity to EGFR TKI therapy in an *EGFR L858R* lung cancer patient whose tumor also harbored a clonal co-occurring truncating *RBM10* mutation (13). Thus, at individual patient resolution across multiple cases RBM10 inactivation was associated with decreased initial EGFR TKI sensitivity, relatively early clinical progression, and decreased pathological tumor response in *EGFR* mutant LA.

To further establish the functional impact of the patient-derived *RBM10* mutations, we engineered the *RBM10 Y36*, RBM10 S167\**, and *RBM10 Q595\** variants and expressed each mutant form in RBM10-deficient, *EGFR* mutant A014 cells (Figure 4F, Supplementary Figure 4D-F, Supplementary Figure 6A; the RBM10 splice site variant in case #4 proved challenging to engineer). In contrast to WT *RBM10*, each *RBM10* mutant failed to rescue the apoptotic phenotype upon osimertinib treatment, as measured by PARP cleavage (Figure 4G). These findings indicate that these *RBM10* mutations present in the human *EGFR* mutant lung cancers are loss-of-function mutations and result in decreased apoptotic response to EGFR TKI treatment.

In summary, these clinical findings mirror the effects of RBM10 deficiency that we observed in the *in vitro* and *in vivo* in the preclinical models, namely diminished initial tumor cell apoptosis and EGFR TKI response, and complement the independent clinical data described above (Figure 4A-B). While larger clinico-genomic cohorts of *EGFR* mutant tumors treated with an EGFR TKI and containing treatment outcome status are lacking, our collective findings provide proof-of-concept of a role for RBM10 deficiency in limiting EGFR TKI response and provide a rationale for further analysis as additional clinical case data become available in the future.

### RBM10 deficiency decreases the Bcl-xS/Bcl-xL ratio to limit the apoptotic response to EGFR TKI therapy

Our studies suggested that loss of *RBM10* can limit EGFR TKI-induced apoptosis rather than increase proliferative capacity in cancer cells. To investigate the mechanism by which RBM10 deficiency decreases treatment-induced apoptosis, we first undertook a global analysis to identify which mRNAs were alternatively spliced in response to RBM10 status using an established independent dataset (36). This dataset was derived from an analysis of RBM10 replete or deficient HEK293 cells. The analysis revealed 412 genes whose differential mRNA splicing was regulated by *RBM10* knockdown (Supplementary Figure 7A, Supplementary Table 1) (36). Biological pathway analysis of these 412 targets using established databases (37–39) revealed a significant enrichment for “cell death pathways” with high statistical significance (*p* < 0.05; 2^nd^ highest rank) (Supplementary Figure 7A). We focused on cell death regulation downstream of RBM10 because we noted that RBM10 deficiency suppressed apoptosis during EGFR TKI treatment in the preclinical models (as shown above).

We next sought to identify the specific apoptosis related genes among these 412 through which RBM10 could function to limit EGFR TKI response. RBM10 is known to regulate mRNA splicing of factors involved in intrinsic (mitochondrial-mediated) apoptosis, such as Bcl-x and Caspase 9 (28). Bcl-x, a member of the Bcl-2 family of proteins, is a mitochondria-associated protein that regulates apoptosis (17). Bcl-x can generate either of two expressed proteins as a result of alternative mRNA splicing: a short pro-apoptotic form, Bcl-xS, and a longer anti-apoptotic form, Bcl-xL (27, 28). We found that *RBM10* knockdown decreased the Bcl-xS-to-Bcl-xL ratio to favor a potential anti-apoptotic phenotype (Supplementary Figure 7B). This relationship with RBM10 status appeared to be specific for Bcl-x, as we noted no other Bcl-2 family genes whose mRNA splicing was significantly affected by *RBM10* knockdown in the global HEK293 cell dataset (Supplementary Table 1) (36). Expression of the truncated isoform of *Caspase 9, Caspase 9b* (also called *Caspase 9s*) that can be regulated by RBM10 in certain contexts and that is anti-apoptotic was not increased by *RBM10* knockdown in this analysis (Supplementary Table 1) (36), again suggesting a specific association between RBM10 status and Bcl-x mRNA splicing.

Given these collective findings, we focused on *Bcl-x* and hypothesized that RBM10 loss could decrease the Bcl-xS-to-Bcl-xL ratio, limiting the apoptotic response to EGFR targeted therapy (Figure 5A). To test this, we performed Bcl-x Q-RT-PCR analysis in H3255 and PC-9 cells that were either RBM10 replete (shScramble) or deplete (*RBM10* KD). We observed a decrease in the Bcl-xS-to-Bcl-xL ratio, with a relative increase in the anti-apoptotic Bcl-xL transcript upon *RBM10* silencing (Figure 5B-E). The decreased Bcl-xS-to-Bcl-xL ratio occurred concomitantly with a decreased apoptotic response upon EGFR TKI treatment in H3255 and PC-9 cells (Figure 5F-G). Genetic rescue of *Bcl-xS* expression in the RBM10-deficient cells restored EGFR TKI-mediated apoptosis to levels comparable to RBM10 replete cells (Figure 5F-I), providing evidence of the functional relevance of *Bcl-xS* expression in the apoptotic response to EGFR TKI treatment.

**Figure 5.**
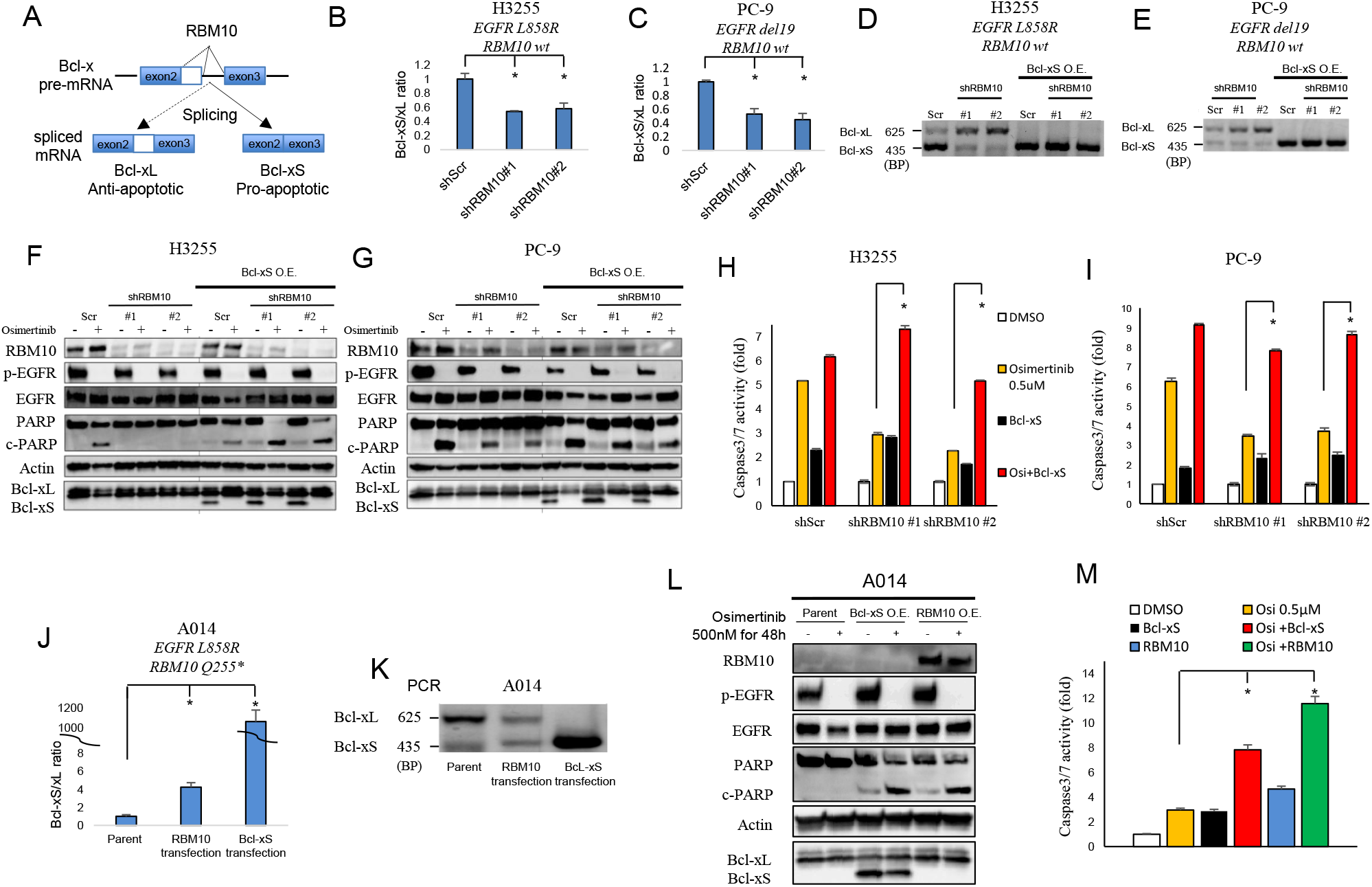
RBM10 deficiency decreases the Bcl-xS/Bcl-xL ratio to limit the apoptotic response to EGFR TKI therapy. **A,** RBM10 regulates *Bcl-x* mRNA splicing into *Bcl-xS* (pro-apoptotic) and *Bcl-xL* (anti-apoptotic) isoforms. **B-C,** Quantitative RT-PCR analysis of Bcl-xS-to-Bcl-xL ratio (mRNA levels) in H3255 (**B**) and PC-9 (**C**) cells expressing *shRBM10* or shScr control. Data are shown as mean ± SEM of the fold change after normalization to housekeeping gene (GAPDH). **D-E,** Conventional PCR analysis using validated primers to detect both Bcl-xL and Bcl-xS isoforms in H3255 and PC-9 cells expressing either shScr control, *shRBM10* with or without genetic rescue of *Bcl-xS*. **F-G,** H3255 and PC-9 (*EGFR L858R, EGFR del19* respectively; *RBM10* WT) cells treated with osimertinib for 48 and 72 hours, which express either *shRBM10* or shScr control paired with or without genetic rescue of Bcl-xS. Cell lysates were harvested, and the indicated proteins were determined by western blot analysis. **H-I,** Caspase-3/7 activity was measured using Caspase-Glo 3/7 assay. Each bar represents the mean ± SEM of the fold change after normalization to DMSO control. **J-K,** Quantitative RT-PCR (**J**) and conventional RT-PCR analysis (**K**) of the Bcl-xS-to-Bcl-xL ratio (mRNA levels) following genetic reconstitution of *RBM10* or *Bcl-xS* in RBM10-deficient A014 cells. **L,** A014 cells (RBM10 deficient) overexpressing *Bcl-xS* or reconstituted with *RBM10* 24h before the osimertinib (500 nM) treatment or DMSO control for 48 hours. Cell lysates were harvested, and the indicated proteins were determined by western blot analysis. **M,** Activity of Caspase-3/7 was measured using Caspase-Glo 3/7 assay. Each bar represents the mean ± SEM of the fold change after normalization to DMSO control. Data represent 3 independent experiments. O.E.: Overexpression. BP; Base Pairs. *; p < 0.05.

We next investigated if exogenous *Bcl-xS* or *RBM10* expression in RBM10-deficient A014 cells could increase the Bcl-xS-to-Bcl-xL ratio to enhance EGFR inhibitor-mediated apoptosis. Indeed, we observed increased PARP cleavage and Caspase 3 and 7 activity in A014 cells with genetic reconstitution of *RBM10* or *Bcl-xS* overexpression when treated with osimertinib (Figure 5J-M). Furthermore, each of the *RBM10* mutants that are present in the clinical cases shown in Figure 4 failed to increase the Bcl-xS-to-Bcl-xL ratio, in contrast to WT RBM10 (Supplementary Figure 6B). These findings further corroborate the loss-of-function effect of these *RBM10* variants. We also studied H1975 (*EGFR L858R/T790M; RBM10* G840fs*7) cells that are RBM10-deficient at baseline and used a lower osimertinib concentration because these cells harbor *EGFR T790M*, which may help to confer relatively increased osimertinib sensitivity (40). Similar to A014 cells, reconstitution of *RBM10* or *Bcl-xS* overexpression in H1975 cells increased the Bcl-xS-to-Bcl-xL ratio and osimertinib-mediated apoptosis (Supplementary Figure 8A-G). The collective data indicate that the decreased EGFR inhibitor-mediated apoptosis that we observed in RBM10-deficient *EGFR* mutant LA is at least in part due to a low Bcl-xS-to-Bcl-xL ratio.

To further understand the cell-based mechanisms of RBM10/Bcl-xS-mediated apoptosis induction, we assessed mitochondrial membrane potential upon EGFR inhibition, as measured by mitochondrial matrix pH using an established ratiometric pH-sensitive probe SypHer-dmito (41, 42). We found that *RBM10* knockdown in PC-9 and H3255 cells exhibited higher mitochondrial matrix pH (higher SypHer-dmito ratio [F470/F430]), indicative of increased mitochondrial membrane potential and decreased apoptosis induction compared with shScramble control (Supplementary Figure 9A-B). In contrast, exogenous expression of *RBM10* or *Bcl-xS* in *RBM10*-deficient H1975 and A014 cells lowered the mitochondrial membrane potential compared to control (Supplementary Figure 9C-D). These findings indicate that RBM10 and Bcl-xS can modulate the mitochondrial apoptotic response to EGFR TKI therapy. While we cannot rule out a role for RBM10 regulated mRNA splicing targets beyond Bcl-x in modulating EGFR TKI sensitivity, an area for future investigation, our data establish an important and previously unknown function for differential *BCL-x* mRNA splicing by RBM10 in this context.

### Resistance caused by RBM10 deficiency in *EGFR* mutant lung cancer can be overcome with Bcl-xL and EGFR inhibitor combination therapy

The Bcl-xS-to-Bcl-xL ratio was decreased upon RBM10 loss, resulting in increased Bcl-xL relative to Bcl-xS expression. Thus, we investigated whether inhibition of the anti-apoptotic isoform of Bcl-x, Bcl-xL, could restore apoptosis in *EGFR* mutant *RBM10* KD cells treated with osimertinib. To test this, we used the BH3 mimetic Bcl-xL inhibitor navitoclax (ABT-263) which binds with high affinity to Bcl-xL (43), in combination with osimertinib and observed decreased viability with enhanced PARP cleavage and Caspase 3 and 7 activity in two independent RBM10-deficient *EGFR* mutant cell lines (A014 – *EGFR L858R; RBM10 Q255\** and H1975 – *EGFR L8585R/T790M; RBM10 G840fs*7*) (Supplementary Figure 4D-G) (Figure 6A-F). Additionally, navitoclax treatment rescued osimertinib-mediated apoptosis in H3255 and PC-9 cells expressing *shRBM10* (Supplementary Figure 10A-D). While navitoclax can target other proteins beyond Bcl-xL, including Bcl-2 (43), we used an independent genetic approach to corroborate the role of Bcl-xL by silencing *Bcl-xL* in RBM10-deficient cell lines (A014 and H1975). Under these conditions, we observed enhanced apoptosis in combination with osimertinib (Figure 6G-H). These findings indicate that pharmacologic or genetic suppression of *Bcl-xL* can overcome EGFR inhibitor resistance in RBM10-deficient *EGFR* mutant LA cells.

**Figure 6.**
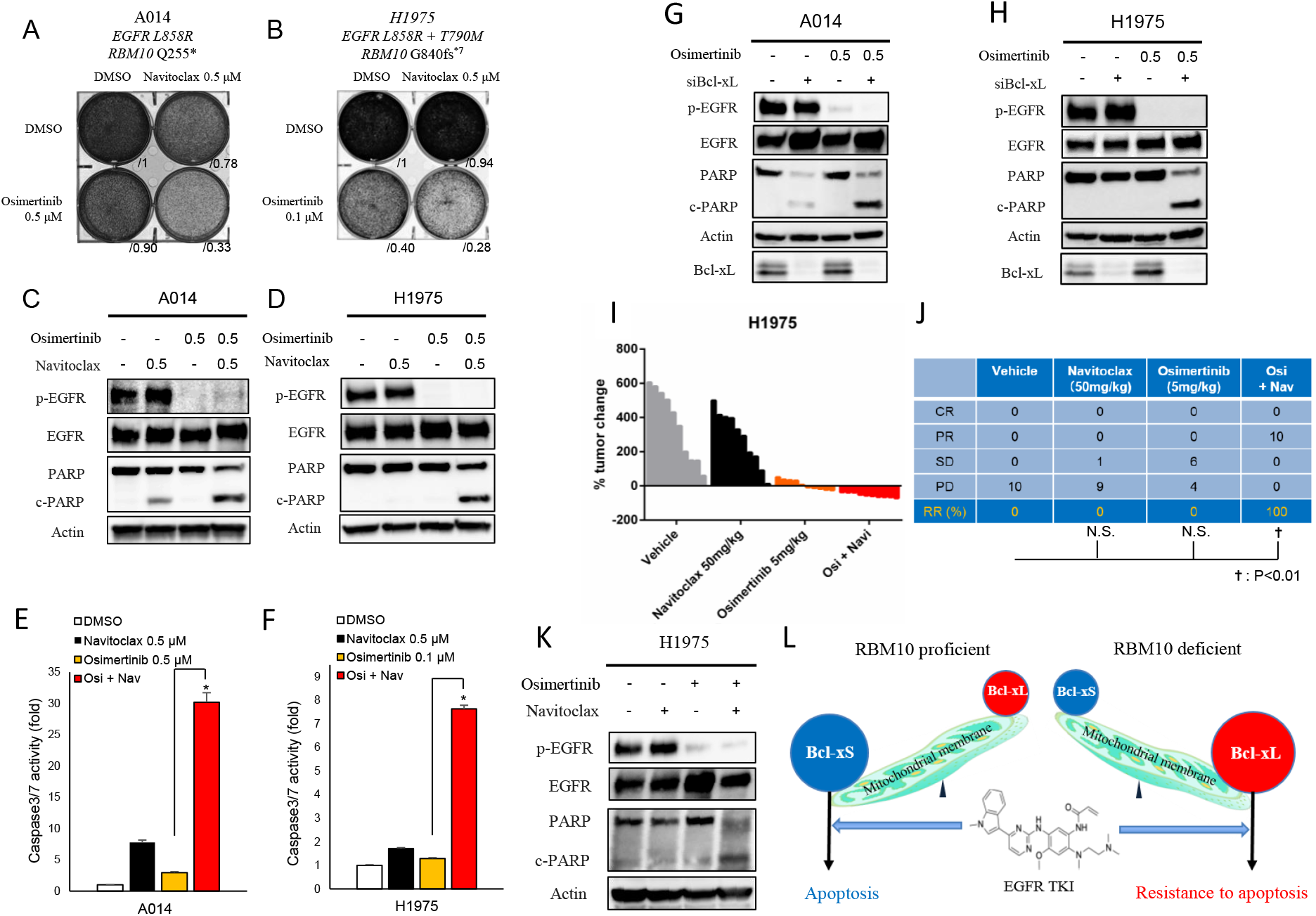
Resistance caused by RBM10 deficiency in *EGFR* mutant lung cancer can be overcome with Bcl-xL and EGFR inhibitor combination therapy. **A-F,** RBM10-defιcient A014 (*EGFR L858R; RBM10 Q255*\*) and H1975 (*EGFR L8585R/T790M; RBM10 G840fs*7*) cells were treated with navitoclax (ABT-263) 500nM alone or in combination with indicated osimertinib concentrations. Crystal violet viability assays (**A, B**) and apoptosis was measured with PARP cleavage and Caspase 3/7 activity (**C-F**). Each bar represents the mean ± SEM of the fold change after normalization to DMSO control (**E, F**). **G-H**, Western blot analysis of *Bcl-xL* knockdown with siRNA in combination with osimertinib 500nM in A014 and H1975 cells. **I-J,** Mice bearing H1975 subcutaneous xenografts were treated with vehicle, navitoclax (50mg/kg), osimertinib (5mg/kg), or combination (navitoclax and osimertinib) therapy for 14 days (n=10 tumors each arm). Percent changes in tumor volume compared to baseline for individual xenografts are shown. Objective tumor response for indicated treatment groups in (**J**). **K,** H1975 xenograft tumor explants were treated with vehicle, navitoclax, osimertinib, or combination (navitoclax and osimertinib, 50mg/kg and 5mg/kg respectively) therapy for 14 days. Tumors were harvested 4 hours after treatment and the indicated protein levels were determined by Western blot analysis. **L,** Proposed model of RBM10 regulating EGFR TKI-induced apoptosis through differential Bcl-x splicing. Data represent 3 independent experiments in data panels. *; p < 0.05.

In order to further validate these therapeutic findings *in vivo*, we treated mice bearing H1975 tumor xenografts with navitoclax, osimertinib, or combination (navitoclax and osimertinib) therapy (Figure 6I-K) (A014 cells did not form tumors *in vivo*). All mice treated with the navitoclax and osimertinib combination achieved an objective response (partial response or complete response by RECIST version1.1 criteria (3)), whereas no objective responses were observed in mice treated with either navitoclax or osimertinib alone (Figure 6I-J). Analysis of the tumor explants revealed an increase in PARP cleavage in H1975 tumors obtained from mice treated with the combination of navitoclax and osimertinib (Figure 6K).

Altogether, these findings indicate that RBM10 deficiency suppresses mitochondrial-mediated apoptosis in response to EGFR inhibition in *EGFR* mutant LA by decreasing the Bcl-xS-to-Bcl-xL ratio (Figure 6L). The EGFR TKI insensitivity induced by RBM10 deficiency can potentially be addressed with combination therapies that target the anti-apoptotic isoform of Bcl-x, Bcl-xL, together with an EGFR TKI.

## Discussion

This study addresses an emerging and important question: what, if any, is the impact of co-occurring genetic alterations in cancers harboring a canonical primary driver mutation, such as mutant *EGFR*? Our data highlight the increasing need to delineate genetic heterogeneity present both within and between *EGFR* mutant tumors and to understand the functional consequences of this genetic heterogeneity to improve clinical outcomes. Our findings provide insight by revealing a previously unappreciated role for co-occurring *RBM10* deficiency in limiting the initial response to EGFR inhibitor treatment in human *EGFR* mutant LA by suppressing tumor cell apoptosis. The role of RBM10 deficiency in limiting tumor cell apoptosis during this early period of initial therapy is distinct from mechanisms that promote the subsequent emergence of acquired resistance after an initial robust tumor response, such as *EGFR T790M* or *MET* kinase amplification, which often drive tumor cell proliferation (8–13). Our data suggest one model for the multi-faceted evolution of resistance such that RBM10 inactivation, for instance via subclonal mutation, may allow for a fraction of cancer cells to avoid apoptosis and persist during initial targeted therapy, resulting in an incomplete tumor response. This is consistent with our observations that tumors with RBM10 deficiency generally are not completely intrinsically resistant and instead show some, suboptimal response while persisting during EGFR TKI treatment. Subsequent frank tumor progression may then occur via the acquisition of other genetic events that further drive acquired resistance, such as drug-resistant secondary mutations in the *EGFR* or the activation of bypass signaling pathways. Thus, RBM10 deficiency may function distinctly from, yet cooperate with, additional molecular events to limit tumor response to EGFR TKI treatment and enable drug-resistant tumor progression that is, over time, lethal in patients.

While other reports indicated recurrent *RBM10* mutations in genetically-unselected LA cases, the relatively low frequency of *EGFR* mutant tumors in these published datasets precluded a subtype-specific analysis (24, 25, 30). *RBM10* truncating mutations were more frequently observed in the *EGFR L858R* subtype compared to tumors that harbored *EGFR exon 19 deletions* (15% vs 3%; p<0.01, Figure 1B). Given the functional role of RBM10 loss in limiting therapeutic response to EGFR inhibitors, our data reveal a potential mechanism to help explain why patients with *EGFR L858R* mutation tumors generally experience worse clinical outcomes and decreased EGFR inhibitor response compared to patients with *EGFR exon 19 deletion* tumors (44). How or why *RBM10* genetic alterations are enriched in the *EGFR L858R* mutant subtype remains unexplained and will be a focus of future investigation.

Our study sheds light on the role of alterations in mRNA splicing factors in cancer pathogenesis. Somatic mutations in genes encoding the spliceosome have been identified in hematopoietic malignancies, including up to 60% of patients with myelodysplastic syndromes (MDS). These mutations commonly occur in *SF3B1* (*Splicing Factor 3b Subunit 1*), *SRSF2* (*Serine/arginine-Rich Splicing Factor 2*), and *U2AF1* (*U2 Small Nuclear RNA Auxiliary Factor 1*) and the genetic data in myelodysplastic syndrome suggests that these alterations are critical to disease pathogenesis (22). Yet, only some of the mutations in the splicing regulators that are recurrently altered in hematopoietic malignancies have been detected in solid tumors to date (23).

Our findings provide an initial example, to our knowledge, of a mechanistic role for splicing factor inactivation (here, RBM10 deficiency) in modulating sensitivity to targeted kinase inhibitor therapy in solid malignancies. RBM10 deficiency does not modulate the oncoprotein target itself (here, mutant EGFR), and instead functions via the differential regulation of the apoptotic machinery in tumor cells. In contrast, truncated forms of mutant BRAF are associated with resistance to BRAF inhibitor treatment as a form of “on-target” therapy resistance in melanoma and lung cancer (45, 46). Whether these truncated mutant BRAF forms arise via alternative mRNA splicing and, if so, the precise splicing factor involved are unresolved questions. Thus, splicing factor deficiency *per se* (here, of RBM10) appears to represent a distinct mechanism of targeted kinase inhibitor therapy resistance from others reported previously (8–12).

Our findings indicate that *RBM10* deficiency in *EGFR* mutant LA tumors decreases the apoptotic response to EGFR inhibitor therapy, leading to tumor progression during EGFR TKI treatment and worse clinical outcomes. RBM10 controls alternative splicing of the apoptosis regulator Bcl-x to generate two isoforms Bcl-xL (anti-apoptotic) and Bcl-xS (pro-apoptotic) (27, 28). Bcl-x is a member of the Bcl-2 family of proteins that exist at the outer mitochondrial membrane. Bcl-2 family proteins regulate mitochondrial outer membrane permeabilization and the release of cytochrome c into the cytoplasm in response to EGFR TKI treatment (47). An alternative splicing event in exon 2 of *Bcl-x* results in two isoforms of *Bcl-x* with antagonistic effects on cell survival: Bcl-xL (long isoform) that is anti-apoptotic, and Bcl-xS (short isoform) that is pro-apoptotic (47). Mechanistically, RBM10 deficiency alters *Bcl-x* splicing to increase the relative abundance of its anti-apoptotic isoform, Bcl-xL to limit apoptosis upon EGFR TKI therapy. This anti-apoptotic molecular effect arising in RBM10-deficient cells can be overcome by Bcl-xL inhibition (pharmacologic or genetic). Our findings indicate that *RBM10* mutation and/or expression status could be a promising biomarker of response to the combination of osimertinib and navitoclax in an ongoing clinical trial (NCT02520778), an area for future investigation that could serve to refine patient selection for treatment and improve clinical outcomes. Beyond Bcl-x, RBM10 may regulate other genes via mRNA alternative splicing that are involved in therapeutic response, an avenue for future study.

In summary, our findings illustrate the utility of understanding the role of co-occurring genetic alterations in oncogene-driven cancers, with translational implications. The impact of RBM10 deficiency in *EGFR* mutant NSCLC established in this study sheds light on the role of tumor genetic heterogeneity in the multi-faceted evolution of therapeutic resistance.

## Methods

### Patients

#### Human EGFR-TKI treated de-identified patient cohort

Fifty-seven non-small cell lung cancer patients with *EGFR* mutations at Catalan Institute of Oncology, Hospital Germans Trias i Pujol (Badalona, Barcelona, Spain) were treated with EGFR-TKIs. CEIM of the Quirónsalud-Catalunya Hospital Group, RD 1090, was granted by the Institutional Review Board (IRB) on 4 December 2015. Informed consent for the analysis was obtained from all patients.

#### UCSF de-identified clinical cases

IRB-approval for the study # 13-12492 was granted by the UCSF IRB. According to federal regulations summarized in 45 CFR 46.102(f), this study does not involve human subjects and did not require further IRB oversight. The requirement for informed consent was waived. Retrospective chart review of patients was carried out by the study investigators to identify patient demographic information, including objective response, PFS, and OS to EGFR TKI therapy. Direct radiographic review was performed by study investigators when possible.

#### Lung adenocarcinoma dataset

Targeted sequencing with a panel of 324 cancer-related genes on lung cancer samples was provided by Foundation Medicine (https://www.foundationmedicine.com/genomic-testing/foundation-one-cdx). The MSK-IMPACT dataset was download from bioportal (https://www.cbioportal.org/study/summary?id=msk_impact_2017). TCGA-luad RNAseq and whole exome sequencing data was obtained from genomic data commons (https://gdc.cancer.gov/) All data was processed using R programming (version 3.6.2). The oncoprint for *EGFR-RBM10* co-mutations was generated using ComplexHeatMap package in R.

#### Global analysis of alternative mRNA splicing regulated by RBM10 in HEK293 cells

Splicing changes induced by RBM10 KD in HEK293 cells were based on RNA-seq data (36). Briefly, the inclusion ration (PSI: percentage splicing in) of each exon in Refseq transcripts as the number of reads supporting inclusion divided by total number of reads supporting inclusion and exclusion of the specific exon. The inclusion ratio between RBM10 and control was computed, and then transformed into Z-value (36). Functional annotation of these differentially spliced genes was carried out using multiple databases (KEGG, BBID and BIOCARTA) (37–39). Adjusted p value cutoff for significant gene sets was set to 0.05. A hypergeometric test was used for pathway enrichment analysis within the algorithm (DAVID 6.8) (37–39).

#### Human tissue and immunohistochemistry

All patient tumor samples analyzed were obtained under IRB-approved protocols with informed consent obtained from each subject under the guidance of the University of California, San Francisco. All relevant ethical regulations were followed.

#### Cell Lines and Culture Reagents

The A014 cell line was kindly provided by Daniel Tan, Cancer Therapeutics Research Laboratory, Division of Medical Sciences, National Cancer Centre Singapore. PC-9, H3255 and H1975 cells were purchased from ATCC. Cells were maintained at 37 °C in a humidified atmosphere at 5% CO_2_ and grown in RPMI medium 1640 supplemented with 10% (v/v %) FBS, 100 IU/mL penicillin, and 100 μg/mL streptomycin.

#### Cell Viability Assay

For crystal violet experiments, cells (3 × 10^5^) were plated in adherent six-well dishes (Corning). After 24 hours, cells were exposed to either vehicle [dimethyl sulfoxide (DMSO)], osimertinib, or navitoclax. Each assay was performed in triplicate, and representative images are shown. Densitometry quantifications were performed with ImageJ software.

#### Cell apoptosis

Cells (3 × 10^3^) were seeded into 96-well, white-walled plates (Corning) and incubated overnight. Cells were subsequently treated with vehicle (DMSO) or the indicated compounds for 48 hours. Cellular apoptosis was analyzed with Caspase-Glo 3/7 assay kits (Promega), which measures Caspase-3/7 activity, in accordance with the manufacturer’s protocol.

#### Western Blot Analysis

Cells (2 × 10^5^) were seeded in six-well plates and rested overnight before drug treatment for 48-72 hours. Whole-cell lysates were prepared by using radio-immunoprecipitation assay buffer (RIPA) [10 mM Tris·Cl (pH 8.0), 1 mM EDTA, 0.1% sodium deoxycholate, 0.1% SDS, 140 mM NaCl] supplemented with protease inhibitor and phosphatase inhibitor (Roche) and clarified by probe sonication and centrifugation (14,000 x g for 15 min. at 4°C). Equal masses of protein (5 ug – 40 ug) were separated by 4—15% of SDS/ PAGE and were transferred onto nitrocellulose membranes (Bio-Rad) for protein blot analysis. Membranes were incubated with primary antibody overnight and were washed and incubated with secondary antibody for 1 hour. Protein bands were visualized using either a fluorescence system (LI-COR) or Amersham ECL chemiluminescent reagent (GE Life Sciences); chemiluminescent signals were visualized with an ImageQuant LAS 4000 instrument (GE Healthcare).

#### Antibodies

Antibodies for Phospho-EGFR (Tyr1068), total-EGFR, phospho-ERK, total-ERK, beta-actin, Bcl-xL, Ki-67 and cleaved-PARP were purchased from Cell Signaling Technology and diluted (according to manufacturer’s recommendations). The RBM10 antibody used for Western blotting was purchased from Santa Cruz Biotechnology and diluted.

#### Orthotopic Lung Xenografts in Immunodeficient Mice

Six-to eight-week-old female SCID CB.17 mice were purchased from Taconic. Specific pathogen-free conditions and facilities were approved by the American Association for Accreditation of Laboratory Animal Care. Surgical procedures were reviewed and approved by the UCSF Institutional Animal Care and Use Committee (IACUC), protocol no. AN107889-03C. To prepare cell suspensions for thoracic injection, cells were mixed with Matrigel matrix (BD Bioscience 356237) on ice yielding a final concentration of 1.0 × 10^5^ cells/μL. Mice were placed in the right lateral decubitus position and anesthetized with 2.5% inhaled isoflurane. A 1-cm surgical incision was made along the posterior medial line of the left thorax, fascia and adipose tissue layers were dissected and retracted to expose the lateral ribs, intercostal space, and the left lung parenchyma. A 30-guage hypodermic needle was used to advance through the intercostal space ~3 mm into the lung tissue. Cells were taken to inject 10 μL (1.0 × 10^6^ cells) of cell suspension directly into the left lung. Visorb 4/0 polyglycolic acid sutures were used for closure of the fascia and skin layer. Mice were observed after the procedure for 1–2 h. For drug treatments, orthotopically implanted tumors were allowed to grow for 1 week before treatment. Mice were treated with either vehicle or osimertinib at the start of week 2 and continued on therapy until the week 9 (post-implantation day 60).

#### In Vivo Bioluminescence Imaging

Mice were imaged at the UCSF Preclinical Therapeutics Core after tumor injection on day 7 with a Xenogen IVIS 100 bioluminescent imaging system. Before imaging, mice were anesthetized with isoflurane and 200 μL of D-Luciferin at a dose of 150 mg/kg body weight was administered by intraperitoneal injection. Weekly monitoring of bioluminescence of the engrafted lung tumors was performed until week 9. Radiance was calculated automatically by using Living Image Software following demarcation of the thoracic cavity (ROI) in the supine position. The radiance unit of photons per s^-1^ /cm2 osr^-1^ is the number of photons per second that leave a square centimeter of tissue and radiate into a solid angle of one steradian (sr).

#### Subcutaneous (s.c.) Tumor Xenograft Studies

All animal experiments were conducted under UCSF IACUC-approved animal protocol no. AN107889-03C. Beas2B, H1975, H3255 and PC-9 tumor xenografts were generated by injection of cells (1 × 10^6^) in a 50/50 mixture for matrigel and PBS into 6-to 8-wk-old female NOD/SCID mice. Mice were randomized to treatment groups once tumors reached an average size of 150 mm^3^. For drug treatments, H1975, H3255 and PC-9 cells were s.c. implanted and allowed to grow to ~200 mm3 (4 wk after implantation). Mice were then treated with vehicle, osimertinib, and/or navitoclax for 15 days.

#### In Vivo Compound Formulation

Osimertinib and navitoclax was obtained from Selleck chemicals and Chemgood, respectively. For all mice studies, osimertinib and navitoclax were administered daily by oral gavage. Osimertinib was dissolved in a v/v% mixture of 7% DMSO, 13% Tween-80, and 80% 5%-glucose, followed by an acid adjustment using an equimolar volume of HCl. Navitoclax was dissolved in 30% PEG400, 60% Phosal 50 PG and 10% ethanol, and vortexed continuously throughout the dosing period.

#### Immunostaining (IHC): Clinical samples and subcutaneous Xenografts

For subcutaneous xenografts studies, mice were sacrificed at the primary endpoint. Tumors were harvested and fixed in 10% neutral buffered formalin for 48 h, embedded in paraffin, and 5-to 10-μm sections were prepared. Sections were subsequently deparaffinized in xylene, rehydrated in a graded ethanol series, and pressure boiled with 1X target retrieval buffer, sodium citrate pH 6.1 (Agilent Dako) for 1 hour. Tissue was treated with 0.3% H_2_O_2_ for 10 minutes, washed with PBS, and incubated with antibodies directed against RBM10 antibody (Bethyl laboratories) overnight. IHC was evaluated by a semiquantitative approach used to assign an H-score (Mean±SEM).

#### Time-lapse imaging

Cells (5 x 10^4) were seeded in a 35mm petri dish containing a 14mm Microwell No.1.5 coverglasss (0.16-0.19mm, MatTek corporation) for adherent live cell imaging. Cells stably expressed SypHer-dmito construct (41). Cells were transfected with either the *RBM10* or *Bcl-xS* constructs for genetic overexpression experiments, or the si*RBM10* variants for genetic knockdown experiments 48 hours before image acquisition. Immediately before imaging, media was replaced with physiological salt solution (PSS) containing 150 mM NaCl, 4 mM KCl, 2 mM CaCl_2_, 1 mM MgCl_2_, 5.6 mM glucose and 25 mM HEPES (pH 7.4) ± 50 mM DCA at 37°C. Images were acquired with CMOS image sensor, ORCA-Flash 4.0 (Hamamatsu, Japan), equipped with 418-442 nm and 450-490 nm excitation filters, and 510-540 nm emission filter. SypHer-dmito fluorescence (F470/F430) ratio was calculated for each cell after subtracting background signal. All images were analyzed with ImageJ software (42). For each cell type, cells with F470/F430 ratio values greater than two s.d. from the mean value were excluded.

Gene knockdown and over-expression assays. All shRNAs for *RBM10* were obtained from Sigma Aldrich (TRCN0000233276 and 0000233277). Sequences for individual shRNAs; TRCN0000233276:

CCGGGACATGGACTACCGTTCATATCTCGAGATATGAACGGTAGTCCATGTCT TTTTG TRCN0000233277:

CCGGCTTCGCCTTCGTCGAGTTTAGCTCGAGCTAAACTCGACGAAGGCGAAG TTTTTG

Stealth RNAi (Dharmacon) Bcl-xL and Stealth RNAi-negative control were used for RNA interference (RNAi) assays as described. The siRNA target sequences were 5’-CUCCUUCGGCGGGGCACUGUGUU-3’ and 5’-CACAGUGCCCCGCCGAAGGAGUU-3’ for Bcl-xL.

#### Plasmid transfections

Bcl-xS and RBM10 were obtained from Addgene (Cambridge, MA). Plasmid transfections required 3 μg/well and were carried out using 0.1% Fugene HD (Promega).

#### PCR

PCR was carried out at 95 °C for 15 s, followed by 27–30 cycles of 95 °C for 15 s, 58.5 °C for 15 s, and 72 °C for 20 s, and finally at 72 °C for 7 min using AmpliTaq Gold PCR Master Mix (Applied Biosystems) with template cDNA equivalent to 15 ng of total RNA and a high concentration (0.75 mM each) of primers. The PCR conditions were semi-quantitative, and no more than 2–5% of input primers were consumed. The primer set for *Bcl-x* simultaneously amplified two alternatively spliced isoforms (*Bcl-xL*; 625 bp, *Bcl-xS*; 435 bp).

*Bcl-x* Fw: AGCTGGAGTCAGTTTAGTGATGTG
*Bcl-x* Rev: TGAAGAGTGAGCCCAGCAGAAC

#### Quantitative RT-PCR assay for gene expression

Cells (3 × 10^5^) were seeded and given 24 hours to adhere at a confluency of 50%. Cells were treated with inhibitors for 24 hours, followed with rapid RNA extraction using a RNeasy Kit (Qiagen). cDNA was prepared from 500 ng total RNA with the SensiFAST cDNA Synthesis Kit (Bioline). Quantitative PCR was performed using The QuantStudio™ 12K Flex Real-Time qPCR system (Applied Biosystems) using Taqman probes (human RBM10: Assay ID, Hs00275935_m1, human Bcl-xL: Assay ID, Hs00236329_m1, Applied Biosystems, human Bcl-xS: Assay ID, Hs00169141_m1,). GAPDH expression was used as an internal control to normalize input cDNA (human GAPDH: Assay ID, Hs02758991_g1, Applied Biosystems). Ratios of the expression level of each gene to that of the reference gene were then calculated.

#### *RBM10* site directed mutagenesis

QuikChange II mutagenesis kit (Agilent) was used to generate all RBM10 mutants. The QuikChange II primer design website was used to guide the generation of mutagenesis primers for the investigated mutants. For transient transfection experiments, Fugene 6 transfection reagent was used following manufacturer protocols. Primer sequences for individual mutants;

*RBM10 Y36*\*: LEFT; cgctatggagccactgacc, RIGHT; tactcgcgaggatatgaacg
*RBM10 S167*\*: LEFT; gtgcaagcacgggaggtt, RIGHT; gcttccatccatcgtgtagc
*RBM10 Q595*\*: LEFT; ggctactactatgacccccaga, RIGHT; cctctccccatcccagtaca
*RBM10 G840fs*7*: LEFT; agagcccaagaggaggaagt, RIGHT; catccgactgccaatgttgt

#### Statistical Methods

One-way ANOVA or student’s t-test was used to calculate P values for comparing 3 or more groups or 2 groups, respectively. Fisher’s exact test was used to compare the response rate between 2 groups. Wilcoxon’s test was used to compare the Kaplan-Meier curves. All statistical analyses were conducted using JMP8 software (SAS Institute Inc.), with *P* < 0.05 considered statistically significant.

## Author contributions

Conception and design: RO, TB

Development of methodology: SN, WW, RO

Acquisition of data: SN, WW, NK, CB, DK, VO, JS, JR, AU

Analysis and interpretation of data: SN, WW, NK, JS

Writing, review and/or revision of the manuscript: SN, WW, RO, TB

Administrative, technical or material support: NK, CB, SA, MM, FH, NC, DT, YK, RO

Study supervision: RR, RO, TB

## Acknowledgements

This research project was conducted with support from the National Institutes of Health (R01CA231300, U54CA224081, R01CA204302, R01CA211052 and R01CA169338) and the Pew and Stewart Foundations to T.G.B. and by NIH/NCI K08CA222625 to R.O.

## Notes

**Potential conflicts of interest:** T.G.B. is an advisor to Novartis, Astrazeneca, Revolution Medicines, Array, Springworks, Strategia, Relay, Jazz, Rain and receives research funding from Novartis and Revolution Medicines.

### Competing Interest Statement

T.G.B. is an advisor to Novartis, Astrazeneca, Revolution Medicines, Array, Springworks, Strategia, Relay, Jazz, Rain and receives research funding from Novartis and Revolution Medicines.

