## Supplementary materials for "Deficiency of the splicing factor RBM10 limits EGFR inhibitor response in *EGFR* mutant lung cancer"

A

*EGFR* mutant lung cancer
MSK impact dataset
(n = 408)

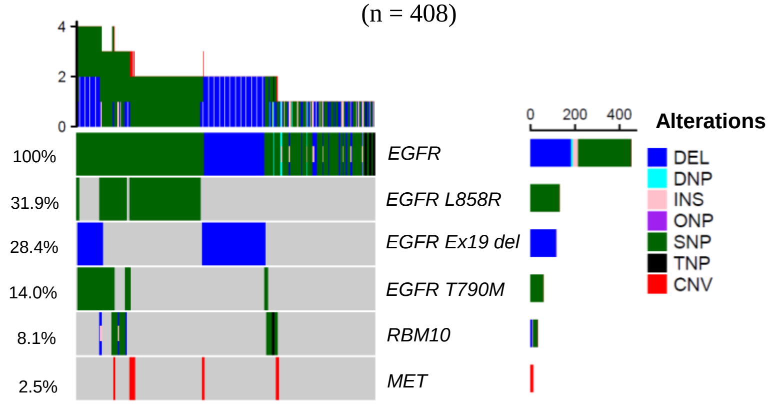


C

*RBM10* and *EGFR* mutation clonality estimation
Foundation Medicine dataset

| | ***EGFR* mutant** | ***RBM10* mutant** |
|---|---|---|
| clonal | 14 | 4 |
| subclonal | 31 | 41 |

p = 0.02 (Fisher exact test)

B

*EGFR* mutant lung cancer
Foundation Medicine dataset
(n = 591)

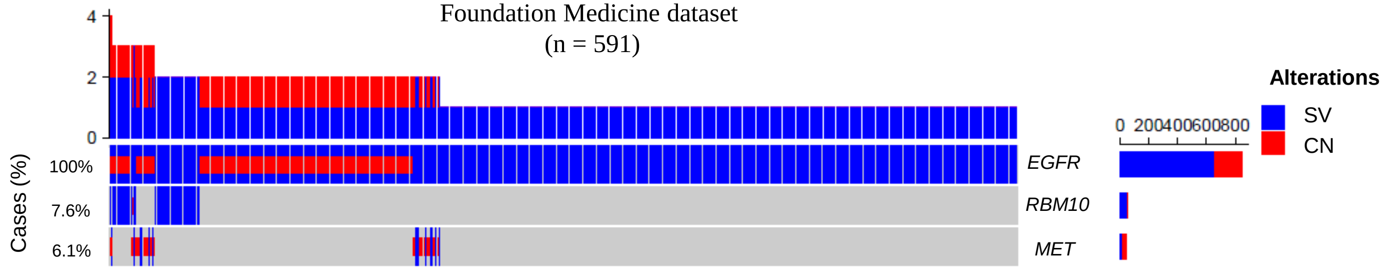

**Supplementary Figure 1. *RBM10* mutations and *MET* gene amplification in MSK-IMAPCT and Foundation Medicine *EGFR* mutant lung adenocarcinoma datasets. A,** Oncoprint of *EGFR* and *EGFR* allelic mutations along with *RBM10* and *MET* alterations in NSCLC. Targeted deep sequencing data from MSK-IMPACT was used to present the concurrent gene alterations in NSCLC. Different colors represent different genetic alterations: DEL=deletion; DNP=Double nucleotide polymorphism; INS=Insertion; SNP=single nucleotide polymorphism; TNP=Triple nucleotide polymorphism; CNV=Copy Number Variation. **B,** Oncoprint of *EGFR-RBM10-MET* gene alterations in lung cancer. Targeted whole exome sequencing data from Foundation Medicine was used to display the pattern of co-alterations. **C,** *RBM10* and *EGFR* mutation clonality estimation in the Foundation Medicine dataset. The distribution (kernel density estimation) of mutational allele frequencies of *RBM10* and *EGFR* mutations were analyzed in this clinical dataset (n=45). A threshold for clonality was set at 0.6 (p = 0.02, Fisher exact test) (30). SV=single nucleotide variant, CN=copy number gain.

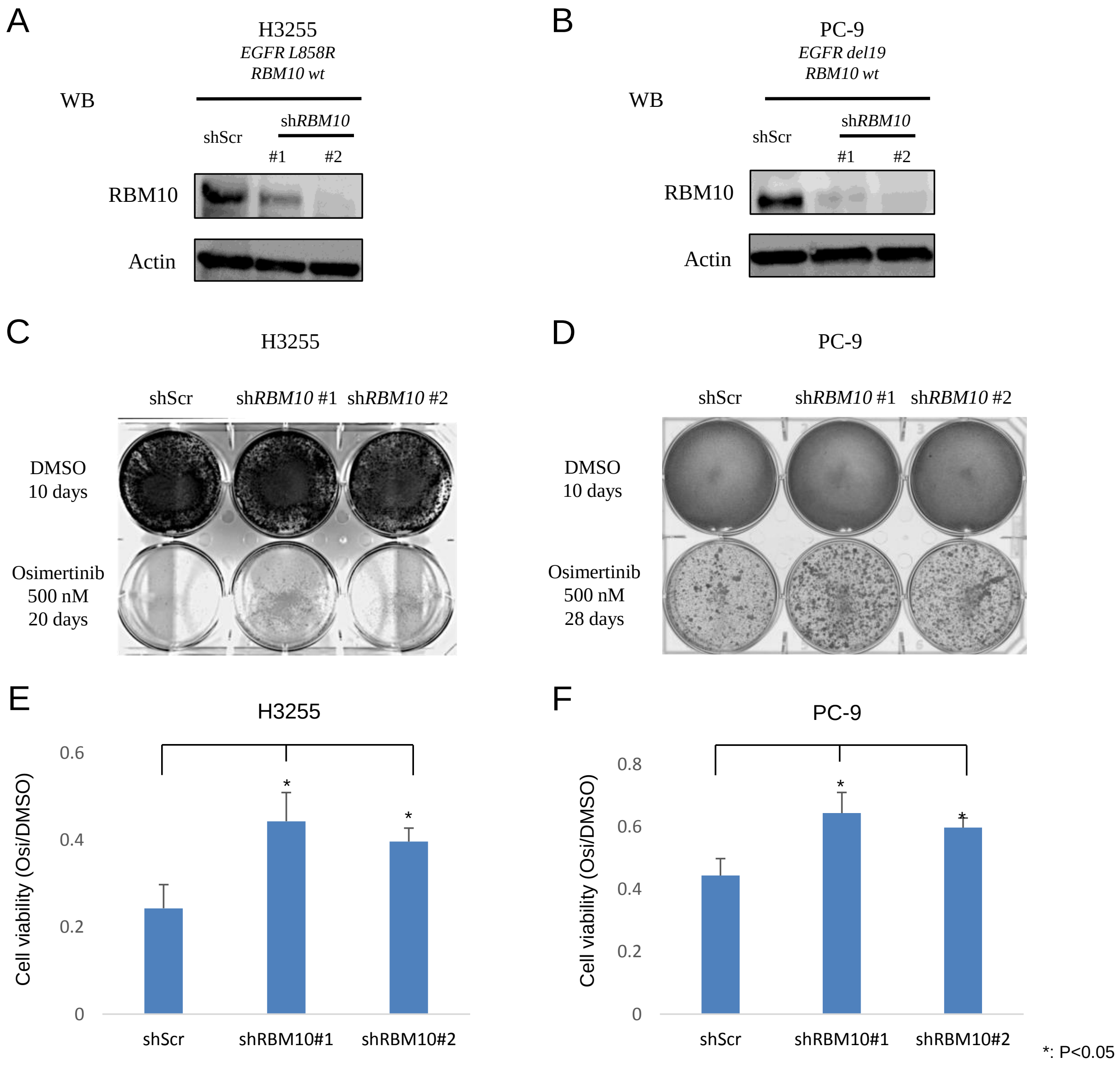
A
H3255
EGFR L858R
RBM10 wt
WB
shScr
shRBM10
#1
#2
RBM10
Actin
B
PC-9
EGFR del19
RBM10 wt
WB
shScr
shRBM10
#1
#2
RBM10
Actin
C
H3255
shScr
shRBM10 #1
shRBM10 #2
DMSO
10 days
Osimertinib
500 nM
20 days
D
PC-9
shScr
shRBM10 #1
shRBM10 #2
DMSO
10 days
Osimertinib
500 nM
28 days
E
H3255
Cell viability (Osi/DMSO)
0
0.2
0.4
0.6
shScr
shRBM10#1
shRBM10#2
*
*
F
PC-9
Cell viability (Osi/DMSO)
0
0.2
0.4
0.6
0.8
shScr
shRBM10#1
shRBM10#2
*
*
*: P<0.05

**Supplementary Figure 2. The effects of osimertinib treatment in RBM10-depleted, *EGFR* mutant cell lines *in vitro*. A-B,** H3255 (**A**) or PC-9 (**B**) cells with shScramble or *RBM10* knockdown confirmed by immunoblot. **C-F,** Crystal violet (CV) cell viability assays demonstrating the effects of osimertinib treatment in each indicated *EGFR* mutant cell line model with *RBM10* knockdown (or control) on growth *in vitro*. Data represent 3 independent experiments (**C, D**). Bars represent the mean ± SD (**E, F**). *; $p < 0.05$.

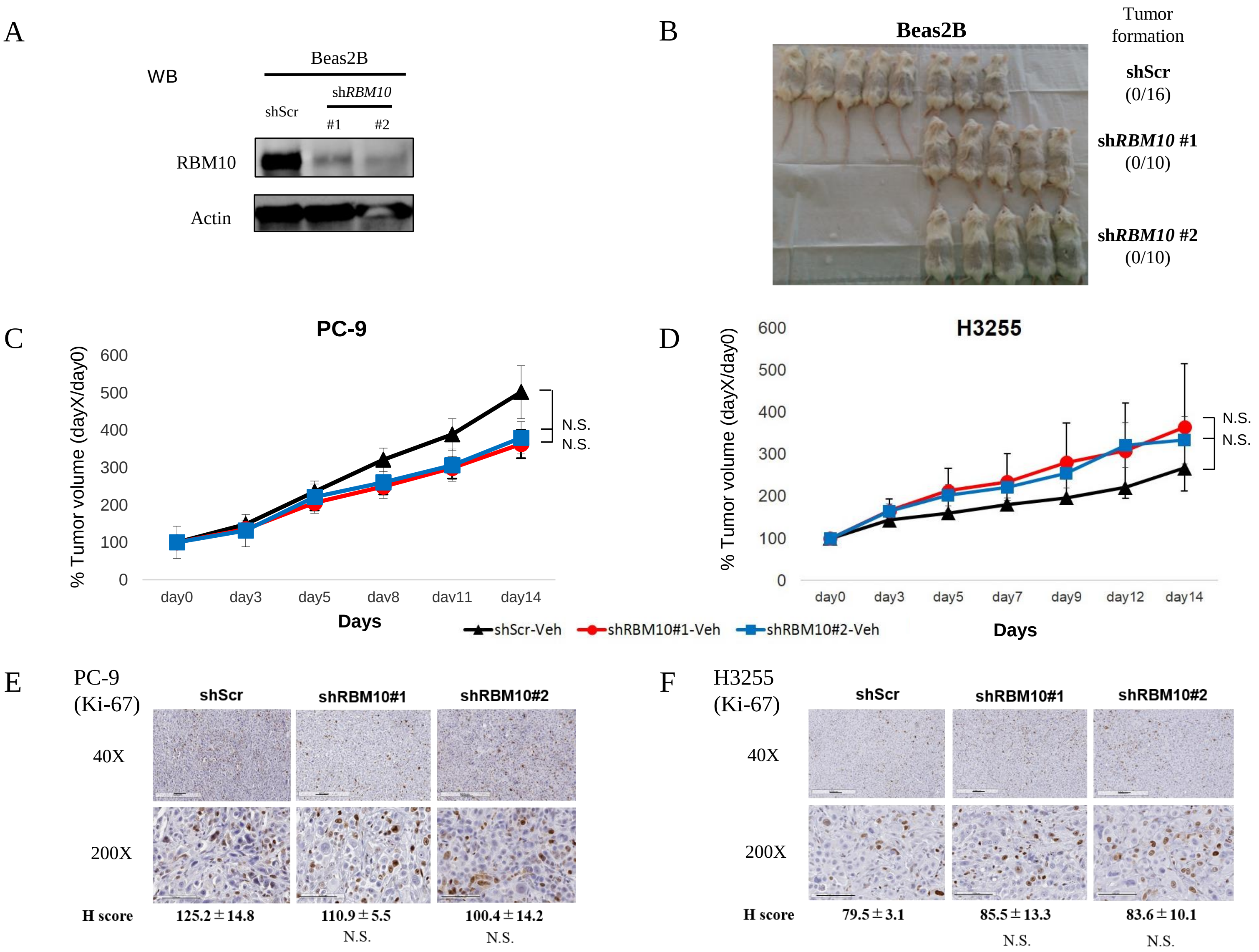
A
WB
Beas2B
shScr
shRBM10
#1
#2
RBM10
Actin
B
Beas2B
Tumor formation
shScr (0/16)
shRBM10 #1 (0/10)
shRBM10 #2 (0/10)
C
PC-9
% Tumor volume (dayX/day0)
Days
N.S.
N.S.
D
H3255
% Tumor volume (dayX/day0)
Days
N.S.
N.S.
shScr-Veh
shRBM10#1-Veh
shRBM10#2-Veh
E
PC-9 (Ki-67)
shScr
shRBM10#1
shRBM10#2
40X
200X
H score
125.2±14.8
110.9±5.5
100.4±14.2
N.S.
N.S.
F
H3255 (Ki-67)
shScr
shRBM10#1
shRBM10#2
40X
200X
H score
79.5±3.1
85.5±13.3
83.6±10.1
N.S.
N.S.

**Supplementary Figure 3. The effects of *RBM10* knockdown on tumorigenesis and tumor growth. A-B,** Beas2B cells with scramble or *RBM10* knockdown were confirmed with the immunoblot (**A**). Those cells were implanted into immunodeficient mice and observed for 60 days (**B**). **C-F,** SCID mice bearing established tumors with PC-9 (**C, E**) or H3255 (**D, F**) cells with shScramble or *RBM10* knockdown were treated with vehicle control once daily for 14 days. Tumor volume was measured using calipers on the indicated days (n=10 tumors each arm). Percent changes in tumor volume compared to baseline for individual xenografts were measured (C-D). Representative IHC analysis of Ki-67 expression with H-score in tumors are shown (E-F). IHC Scale bars; 100 μm and magnification at 40X and 200X. The mean ± SEM for each group are shown. N.S.; not significant.

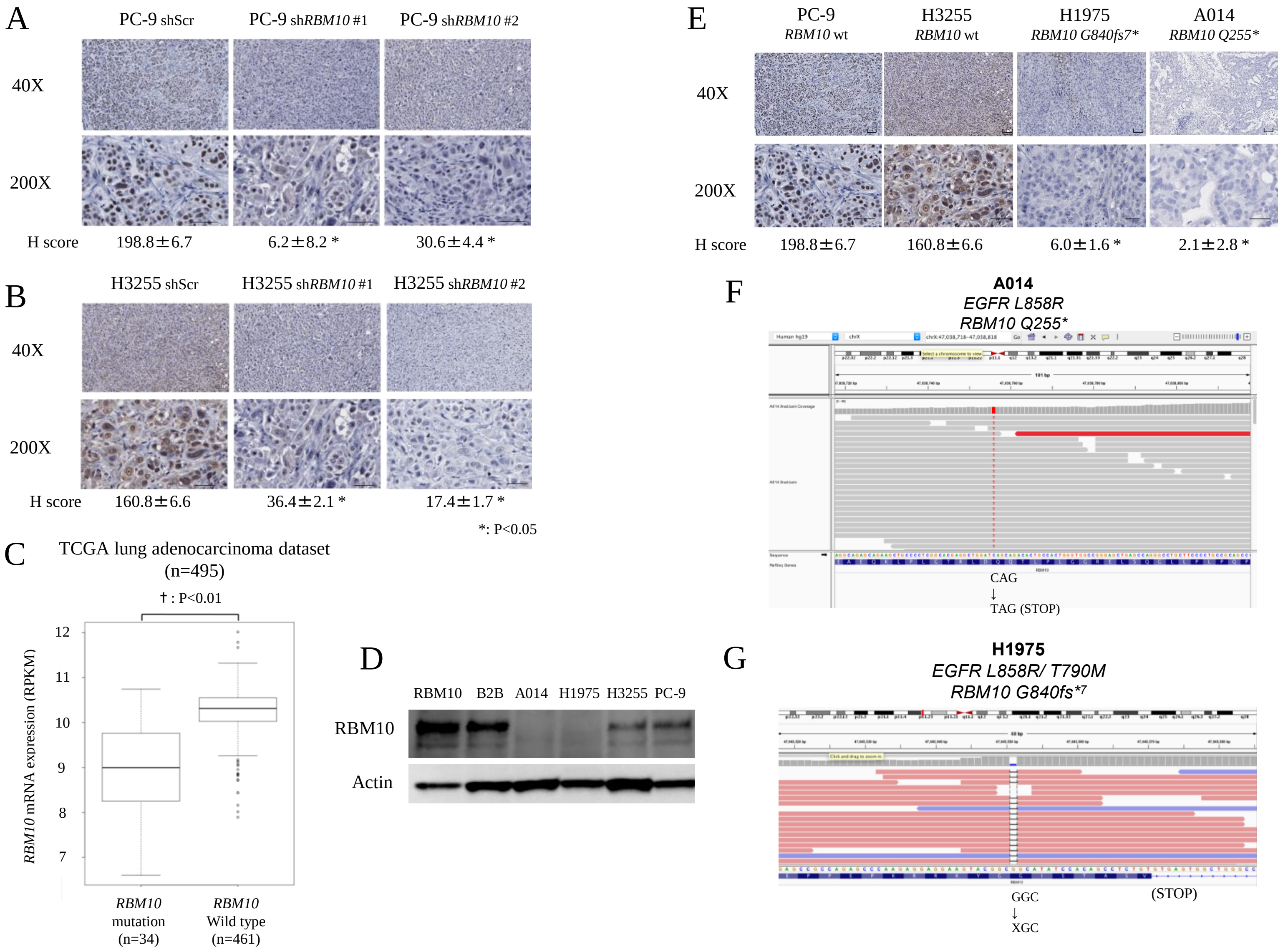
A
PC-9 shScr
PC-9 shRBM10 #1
PC-9 shRBM10 #2
40X
200X
H score
198.8±6.7
6.2±8.2 *
30.6±4.4 *
B
H3255 shScr
H3255 shRBM10 #1
H3255 shRBM10 #2
40X
200X
H score
160.8±6.6
36.4±2.1 *
17.4±1.7 *
*: P<0.05
C
TCGA lung adenocarcinoma dataset
(n=495)
†: P<0.01
RBM10 mRNA expression (RPKM)
12
11
10
9
8
7
RBM10 mutation (n=34)
RBM10 Wild type (n=461)
D
RBM10 B2B A014 H1975 H3255 PC-9
RBM10
Actin
E
PC-9 RBM10 wt
H3255 RBM10 wt
H1975 RBM10 G840fs7*
A014 RBM10 Q255*
40X
200X
H score
198.8±6.7
160.8±6.6
6.0±1.6 *
2.1±2.8 *
F
A014
EGFR L858R
RBM10 Q255*
CAG
↓
TAG (STOP)
G
H1975
EGFR L858R/ T790M
RBM10 G840fs*7
GGC
↓
XGC
(STOP)

**Supplementary Figure 4. RBM10 expression in tumor xenograft and clinical samples.** **A-B,** Representative IHC images of RBM10 in tumors harvested from H3255 and PC-9 subcutaneous tumor xenografts expressing either shScr or sh*RBM10*. **C,** *RBM10* mRNA expression levels in TCGA lung adenocarcinoma samples. *RBM10* mutant and WT samples were stratified using whole exome mutation data and the *RBM10* mRNA expression levels were obtained from corresponding RNAseq data. The boxplot with whisker plot shows the minimum, median, maximum, first and third quantile expression levels. The Y-axis shows the normalized gene expression (RPKM; Reads Per Kilobase Million). †: $p < 0.01$. **D,** RBM10 protein expression in a panel of cell lines. **E,** Representative IHC analysis of RBM10 expression and H-score in tumors harvested from subcutaneous tumor xenografts of H3255 and PC-9 (*RBM10* WT), and H1975 (*RBM10 G840fs**) and A014 (*RBM10 Q255**) cells. **F-G,** Whole exome sequencing findings for A014 (*RBM10 Q255**) and H1975 (*RBM10 G840fs**) indicating the C to T change and frameshift mutation, respectively. Scale bars; 100 μm and magnification at 40X and 200X. *; $p < 0.05$.

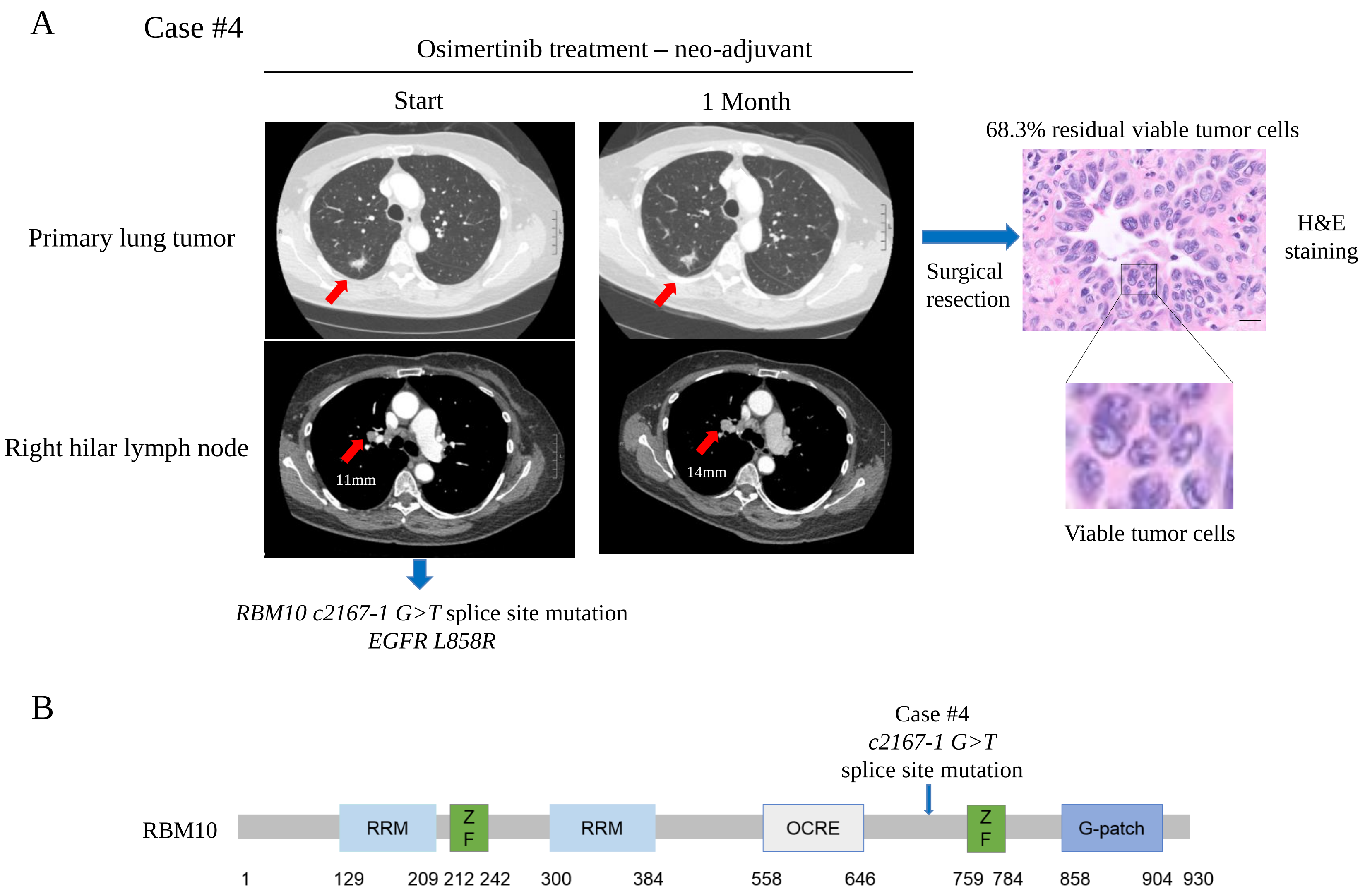
A
Case #4
Osimertinib treatment – neo-adjuvant
Start
1 Month
Primary lung tumor
Right hilar lymph node
11mm
14mm
Surgical resection
68.3% residual viable tumor cells
H&E staining
Viable tumor cells
*RBM10 c2167-1 G>T* splice site mutation
*EGFR L858R*
B
Case #4
*c2167-1 G>T*
splice site mutation
RBM10
RRM
ZF
RRM
OCRE
ZF
G-patch
1
129
209 212 242
300
384
558
646
759 784
858
904 930

**Supplementary Figure 5. *RBM10* mutation is associated with suboptimal EGFR TKI response in *EGFR* mutant NSCLC. A,** Case #4 was a patient enrolled in a neo-adjuvant osimertinib clinical trial and found to harbor co-occurring *EGFR L858R* and *RBM10 c2167-1 G>T* splice site mutations. Following one month of osimertinib treatment, radiographic measurements indicated no substantial regression of the primary lung tumor or the enlarged hilar lymph node. Following two months of osimertinib treatment, pathologic evaluation of the resected tumor specimen showed 68.3% residual tumor cell viability by H&E staining (200X magnification). Scale bar; 100 µm. **B,** The position of the *RBM10 c2167-1 G>T* splice site mutation is shown.

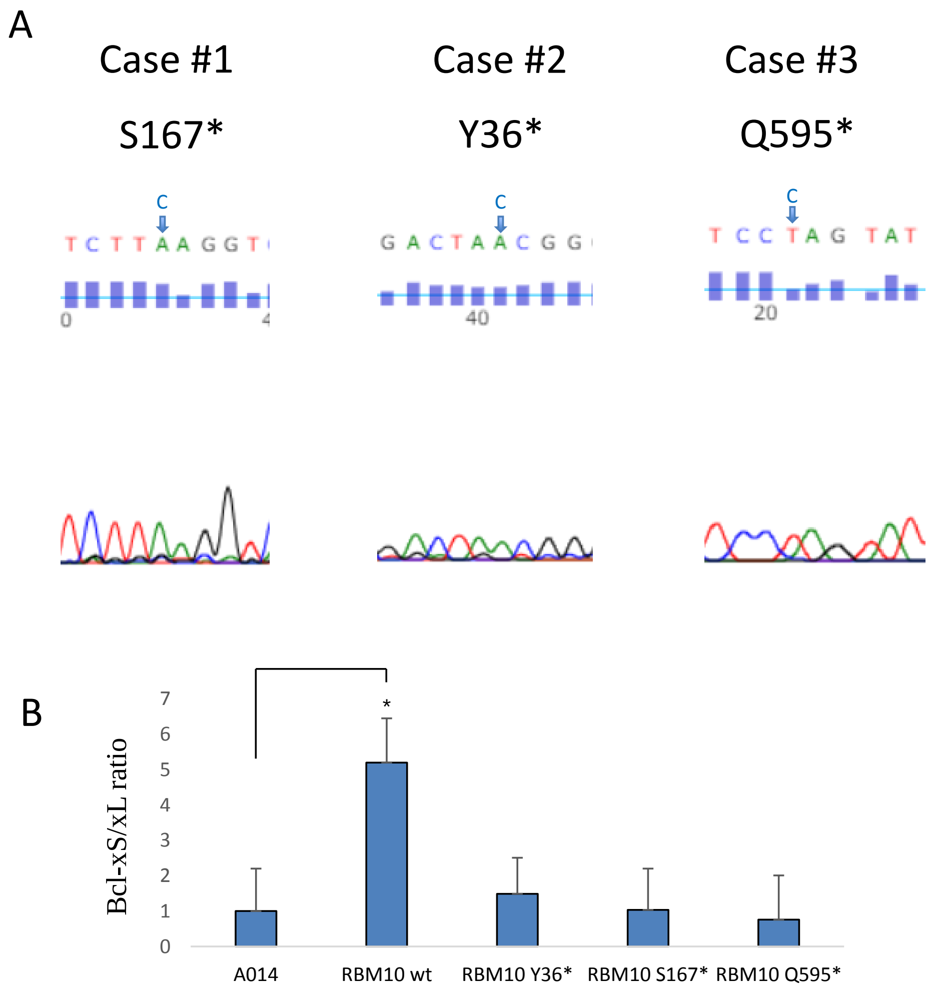
A
Case #1
S167*
Case #2
Y36*
Case #3
Q595*
B
Bcl-xS/xL ratio
A014
RBM10 wt
RBM10 Y36*
RBM10 S167*
RBM10 Q595*

**Supplementary Figure 6. Effect of clinically annotated *RBM10* mutations on the Bcl-xS/Bcl-xL ratio. A,** RT-PCR analysis of A014 (*RBM10 Q255**) cells transfected with *RBM10* mutations (*Y36**, *S167** and *Q595**) constructs. Mutant sequence is shown, with reference nucleotide in blue. **B,** Quantitative RT-PCR analysis of the Bcl-xS-to-Bcl-xL ratio (mRNA levels) in A014 cells with *RBM10* mutations (*Y36**, *S167** and *Q595**) and WT *RBM10*. Data represent 3 independent experiments.

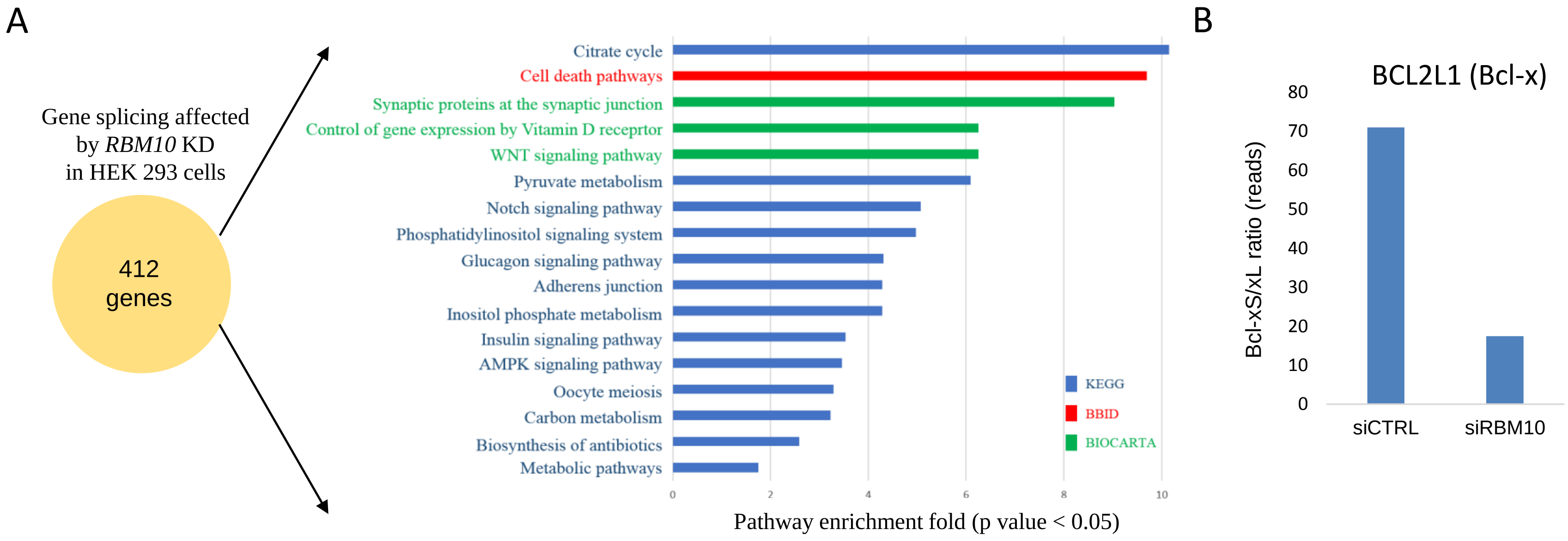


**Supplementary Figure 7. Global analysis of alternative mRNA splicing regulated by *RBM10* in HEK293 cells. A,** mRNA splicing of 412 genes is differentially regulated by *RBM10* knockdown (si*RBM10* for 72h) in HEK293 cells. Functional annotation of these differentially spliced genes was carried out using multiple databases (KEGG, BBID and BIOCARTA). The data are represented as a bar diagram indicating probability of fold pathway enrichment. Adjusted p value cutoff for significant gene sets was set to 0.05. Hypergeometric test was used for pathway enrichment analysis within the algorithm (DAVID 6.8). **B,** A trend towards a decrease in the Bcl-xS-to-Bcl-xL ratio, derived from RNA deep sequencing reads, in HEK293 cells with *RBM10* knockdown (si*RBM10*) compared to control (siCTRL) was observed ($p = 0.06$).

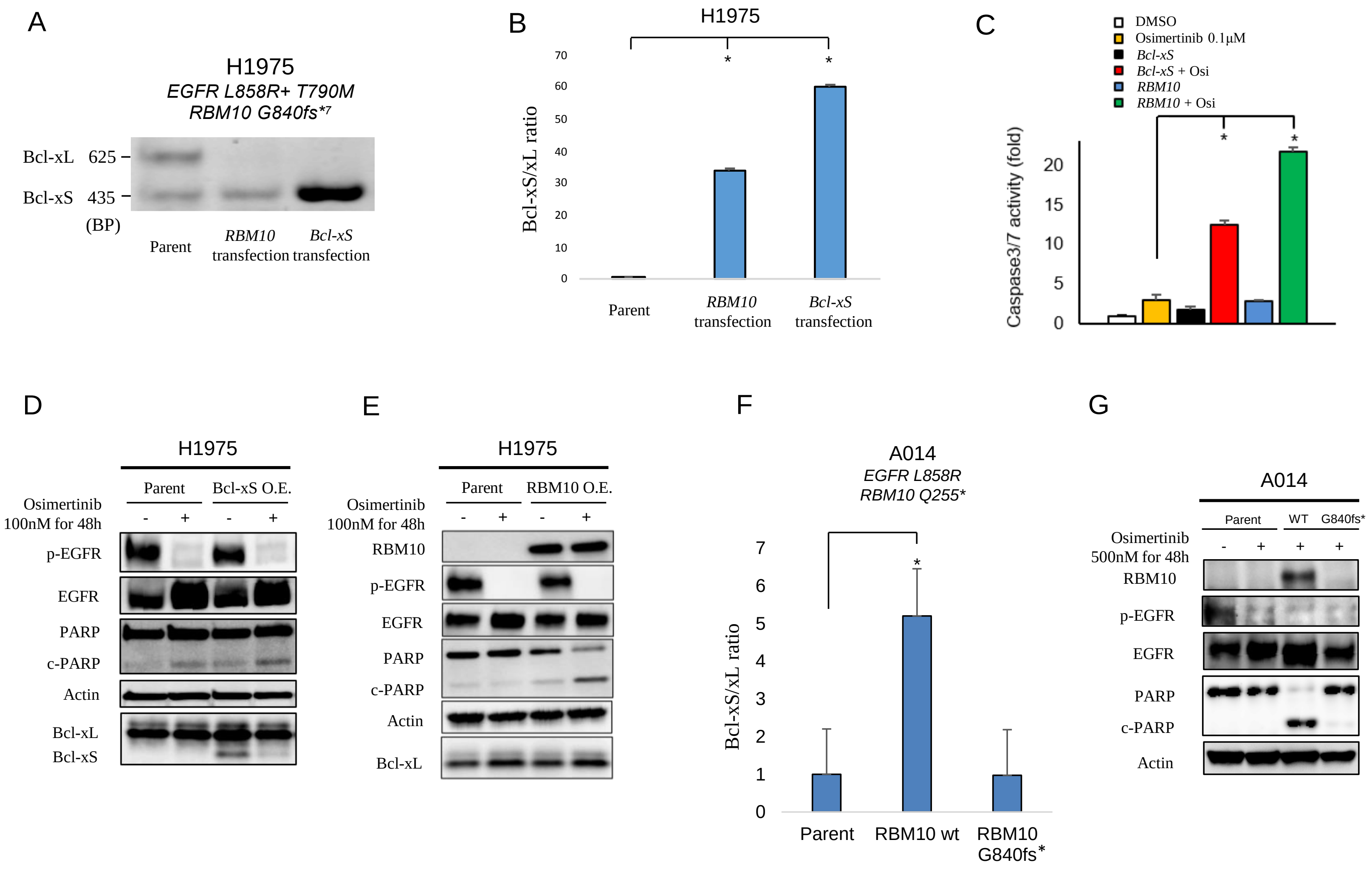
A
H1975
EGFR L858R+ T790M
RBM10 G840fs*7
Bcl-xL 625
Bcl-xS 435
(BP)
Parent
RBM10 transfection
Bcl-xS transfection
B
H1975
Bcl-xS/xL ratio
Parent
RBM10 transfection
Bcl-xS transfection
C
DMSO
Osimertinib 0.1μM
Bcl-xS
Bcl-xS + Osi
RBM10
RBM10 + Osi
Caspase3/7 activity (fold)
D
H1975
Parent
Bcl-xS O.E.
Osimertinib 100nM for 48h
p-EGFR
EGFR
PARP
c-PARP
Actin
Bcl-xL
Bcl-xS
E
H1975
Parent
RBM10 O.E.
Osimertinib 100nM for 48h
RBM10
p-EGFR
EGFR
PARP
c-PARP
Actin
Bcl-xL
F
A014
EGFR L858R
RBM10 Q255*
Bcl-xS/xL ratio
Parent
RBM10 wt
RBM10 G840fs*
G
A014
Parent
WT
G840fs*
Osimertinib 500nM for 48h
RBM10
p-EGFR
EGFR
PARP
c-PARP
Actin

**Supplementary Figure 8. Phenotype of the RBM10-deficient H1975 cell line. A,** Conventional PCR analysis using validated primers to detect both *Bcl-xL* and *Bcl-xS* isoforms in H1975 cells with genetic rescue of *RBM10* or *Bcl-xS* compared to parental control is shown. **B,** Quantitative RT-PCR analysis of the Bcl-xS-to-Bcl-xL ratio (mRNA levels) in H1975 cells with genetic rescue of *RBM10* or *Bcl-xS* compared to parental control cells. **C, D, E,** H1975 cells (RBM10-deficient) overexpressing *Bcl-xS* or reconstituted with *RBM10* 24 hours before treatment with osimertinib (100 nM) or DMSO control. The activity of Caspase-3/7 was measured using Caspase-Glo 3/7 assay. Each bar represents the mean ± SEM of the fold change after normalization to DMSO control (**C**). Cell lysates were harvested, and the indicated proteins were determined by western blot analysis (**D, E**). **F,** Quantitative RT-PCR analysis of the Bcl-xS-to-Bcl-xL ratio (mRNA levels) in A014 (*RBM10 Q255**) cells expressing *RBM10* WT or the *RBM10* mutation native to H1975 (*G840fs*[7]*). **G,** Immunoblot analysis of A014 cells transfected with expression constructs containing *RBM10* WT or the H1975 *RBM10* variant (*G840fs*[7]*). Cells were treated with osimertinib (500 nM) or DMSO for 48 hours and western blot analysis was performed on cellular extracts. Data represent 3 independent experiments. O.E.: Overexpression. BP; Base Pairs. *; $p < 0.05$.

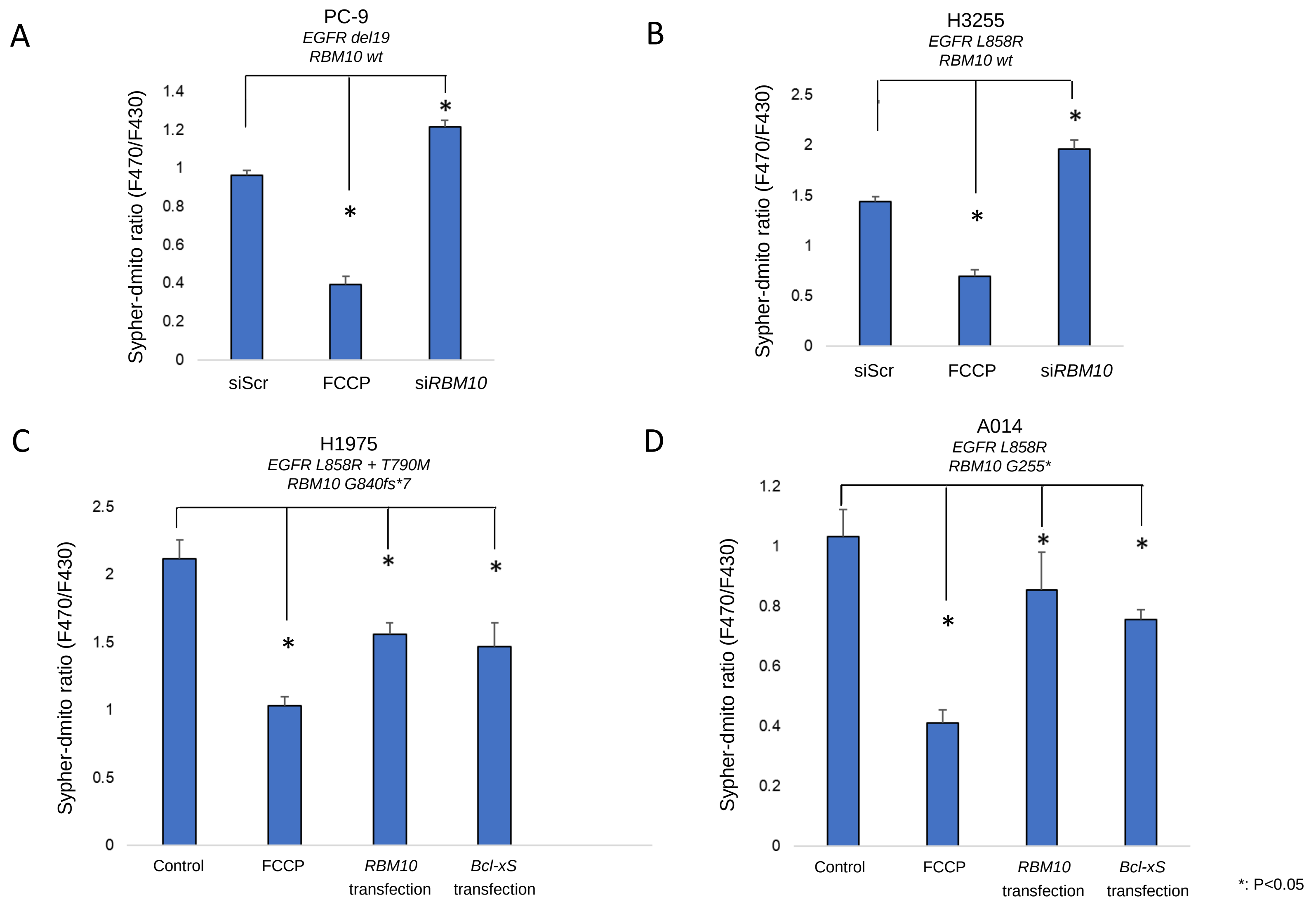
A
PC-9
EGFR del19
RBM10 wt
Sypher-dmito ratio (F470/F430)
1.4
1.2
1
0.8
0.6
0.4
0.2
0
*
*
siScr
FCCP
siRBM10
B
H3255
EGFR L858R
RBM10 wt
Sypher-dmito ratio (F470/F430)
2.5
2
1.5
1
0.5
0
*
*
siScr
FCCP
siRBM10
C
H1975
EGFR L858R + T790M
RBM10 G840fs*7
Sypher-dmito ratio (F470/F430)
2.5
2
1.5
1
0.5
0
*
*
*
Control
FCCP
RBM10 transfection
Bcl-xS transfection
D
A014
EGFR L858R
RBM10 G255*
Sypher-dmito ratio (F470/F430)
1.2
1
0.8
0.6
0.4
0.2
0
*
*
*
Control
FCCP
RBM10 transfection
Bcl-xS transfection
*: P<0.05

**Supplementary Figure 9. Change of mitochondrial membrane potential in response to RBM10 differential expression. A-D,** Cells were stably expressed with SypHer-dmito construct. Cells were treated with uncoupling reagent carbonyl cyanide 4-(trifluoromethoxy) phenylhydrazone (FCCP) as a positive control. SiScramble or si*RBM10* conditions are shown for PC-9 (**A**) and H3255 (**B**) cell lines. RBM10 or Bcl-xS expression constructs for H1975 (**C**) and A014 (**D**) cell lines were transfected 48 hours before image acquisition. SypHer-dmito fluorescence (F470/F430) ratio was calculated for each cell and analyzed with ImageJ software. Each bar represents the mean ± SEM. *: $p < 0.05$ compared to control of each cell.

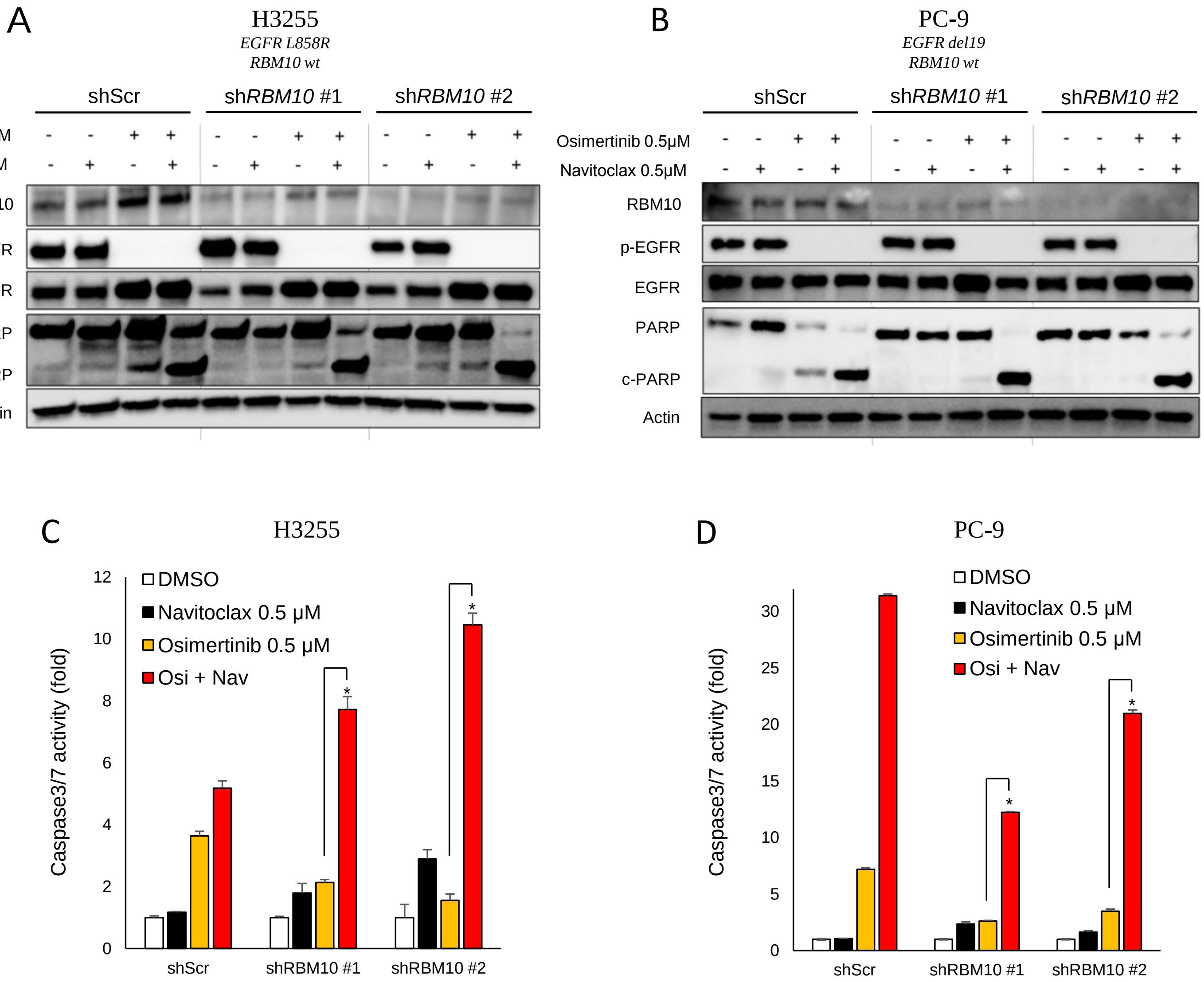
A
H3255
EGFR L858R
RBM10 wt
shScr
shRBM10 #1
shRBM10 #2
Osimertinib 0.5μM
Navitoclax 0.5μM
RBM10
p-EGFR
EGFR
PARP
c-PARP
Actin
B
PC-9
EGFR del19
RBM10 wt
C
H3255
DMSO
Navitoclax 0.5 μM
Osimertinib 0.5 μM
Osi + Nav
Caspase3/7 activity (fold)
D
PC-9

**Supplementary Figure 10. Navitoclax effects in RBM10-depleted cell lines. A-D,** Navitoclax (ABT-263) 500 nM, in combination with osimertinib 500 nM treatment in H3255 ($EGFR^{L858R}$; $RBM10^{\mathrm{WT}}$), PC-9 ($EGFR^{del19}$; $RBM10^{\mathrm{WT}}$) models with or without *RBM10* knockdown. Western blot analysis (**A, B**) and Caspase 3/7 activity (**C, D**) are shown. Each bar represents the mean ± SEM of the fold change after normalization to DMSO control (**C, D**). *; $p < 0.05$. Data represent 3 independent experiments.

**Supplementary Table 1. Differentially spliced exons upon *RBM10* knockdown.** RefSeq exons with significant Percentage Splicing In (PSI) changes (FDR ≤ 5%, ΔPSI ≥ 0.1) after *RBM10* KD in HEK-293 cells (36). Significant Z-values are shown in bold. PSI values in HEK are median between replicates.
